# Transcriptome-wide RNA accessibility mapping reveals structured RNA elements and pervasive conformational rearrangements under stress

**DOI:** 10.1101/2025.06.05.658101

**Authors:** Kelsey Farenhem, Troy W. Whitfield, Alina Chouloute, Ankur Jain

**Affiliations:** Whitehead Institute for Biomedical Research, 455 Main Street, Cambridge, MA 02142, USA; Department of Biology, Massachusetts Institute of Technology, 31 Ames Street, Cambridge, MA 02139, USA

## Abstract

RNA structure plays a central role in post-transcriptional gene regulation, modulating RNA stability, translation, and interactions with RNA-binding proteins (RBPs). However, capturing RNA conformations at scale remains challenging. Here, we introduce DMS-TRAM-seq (Dimethyl Sulfate–Transcriptome-wide RNA Accessibility Mapping by sequencing), which probes RNA structure across nearly the entire transcriptome. Using DMS-TRAM-seq, we generated secondary structure predictions for over 9,000 human transcripts, including hundreds of non-coding RNAs, and identified more than 700 previously unannotated, high-confidence structured elements. Importantly, the enhanced coverage provided by DMS-TRAM-seq enabled comparative analyses, revealing RNAs that undergo structural rearrangements in response to cellular perturbations. Integration with RBP motifs and ribosome profiling uncovered altered RNA–RBP interactions during oxidative stress and showed that translation inhibition broadly drives RNAs toward their thermodynamically favored conformations. DMS-TRAM-seq enables interrogation of the RNA structurome and its plasticity at an unprecedented scale, opening new directions for elucidating the structural basis of RNA regulation.

## INTRODUCTION

Besides encoding genetic information, RNA also performs diverse regulatory and catalytic functions in the cell^1^. This versatility of RNA functions stems from its ability to fold into intricate three-dimensional structures and engage in dynamic interactions with proteins, metabolites, and other RNAs. RNA conformations can change in response to physiological or environmental cues^2–4^, influencing splicing^5^, stability^6,7^, localization^8–10^, and translation^11–13^. While individual RNA structures have been linked to specific regulatory outcomes, many such elements likely remain undiscovered. A comprehensive understanding of the RNA structurome – the complete set of RNA structures across the transcriptome – and how it responds to cellular cues is essential for elucidating the principles of RNA-mediated regulation. However, capturing RNA structural information at the required scale and resolution remains technically challenging, underscoring the need for new methods to probe the RNA structurome.

Over the last two decades, there have been monumental advances in RNA sequencing and associated high-throughput technologies^14^. Conventional RNA sequencing experiments provide information on transcript abundance and sequence-level changes, such as those introduced by alternative splicing. Numerous specialized sequencing-based approaches have also been developed to study specific facets of post-transcriptional RNA regulation. For example, ribosome profiling^15,16^ provides information on a transcript’s translational status, while CLIP-seq^17–19^ captures binding sites of specific RNA-binding proteins (RBPs). Likewise, various techniques have also been developed to examine post-transcriptional base modifications^20,21^. These methods, however, do not provide information on transcripts’ conformational states, underscoring a critical gap in our ability to explore how RNA structure influences cellular function.

High-resolution approaches such as X-ray crystallography^22^ and electron microscopy^23,24^ have provided detailed structural insights into individual RNAs. However, these methods are inherently low-throughput and often require that the RNA adopts a single, stable conformation. As a result, they are typically limited to short, well-folded RNA domains and are not suitable for systematically discovering new folded elements across the transcriptome.

Chemical probing methods, such as in-line probing^25,26^, DMS (dimethyl sulfate)^27,28^, glyoxal^29,30^, and SHAPE (selective 2′-hydroxyl acylation analyzed by primer extension) reagents^31–33^, offer scalable strategies for investigating RNA structure by leveraging reactivity dependent on either base-pairing or backbone flexibility. For example, in-line probing exploits the intrinsic propensity of RNA to undergo spontaneous hydrolysis in alkaline conditions at structurally flexible sites^25^. DMS selectively modifies unpaired A and C bases at their Watson-Crick faces, as well as G bases, albeit at a position not involved in base pairing^27^, while SHAPE reagents modify the 2’-hydroxyl group of RNA nucleotides with flexible backbones, serving as indicators of unpaired regions^4,34^. These techniques have also been coupled with high-throughput sequencing, where nucleotide-resolution reactivity data is read out either by early termination of reverse transcription at modified sites or by incorporation of mismatches during reverse transcription, a strategy known as mutational profiling (MaP). Both DMS mutational profiling by sequencing (DMS-MaPseq)^35^ and SHAPE-MaP^36^ have been used to predict secondary structures of various RNAs of interest, including viral genomes^37^ and mitochondrial RNAs^38^. Although, in principle, these approaches could facilitate RNA structural analysis at a transcriptome-wide scale, experimental realization of such scale has been challenging. Technical limitations, such as difficulty in obtaining large amounts of high-quality RNA after chemical probing, the challenges in using random priming with reverse transcriptases compatible with mutational profiling, and the high degree of duplication in the sequencing libraries, have collectively led to insufficient coverage for robust transcriptome-wide investigations. Consequently, prior studies using mutational profiling have typically been restricted to a few hundred RNA regions with adequate data quality^35,38–41^.

Here, we introduce DMS transcriptome-wide RNA accessibility mapping by sequencing (DMS-TRAM-seq), a mutational profiling approach that builds upon DMS-MaPseq^35^ and provides quantitative, nucleotide-resolution information on RNA accessibility *in vivo* for nearly all expressed transcripts (Figure 1). These data serve as *in cellulo* constraints for transcriptome-scale secondary structure prediction, which we make available through an accessible web portal (humanRNAmap.wi.mit.edu). By systematically analyzing the mutational profiles, we generated secondary structure predictions for over 9,000 human RNAs and uncovered hundreds of novel high-confidence structured elements. Integration with ribosome profiling further showed how translation shapes RNA structure, including a global tendency for RNAs to adopt thermodynamically favored conformations upon inhibition of translation initiation.

**Figure 1.**
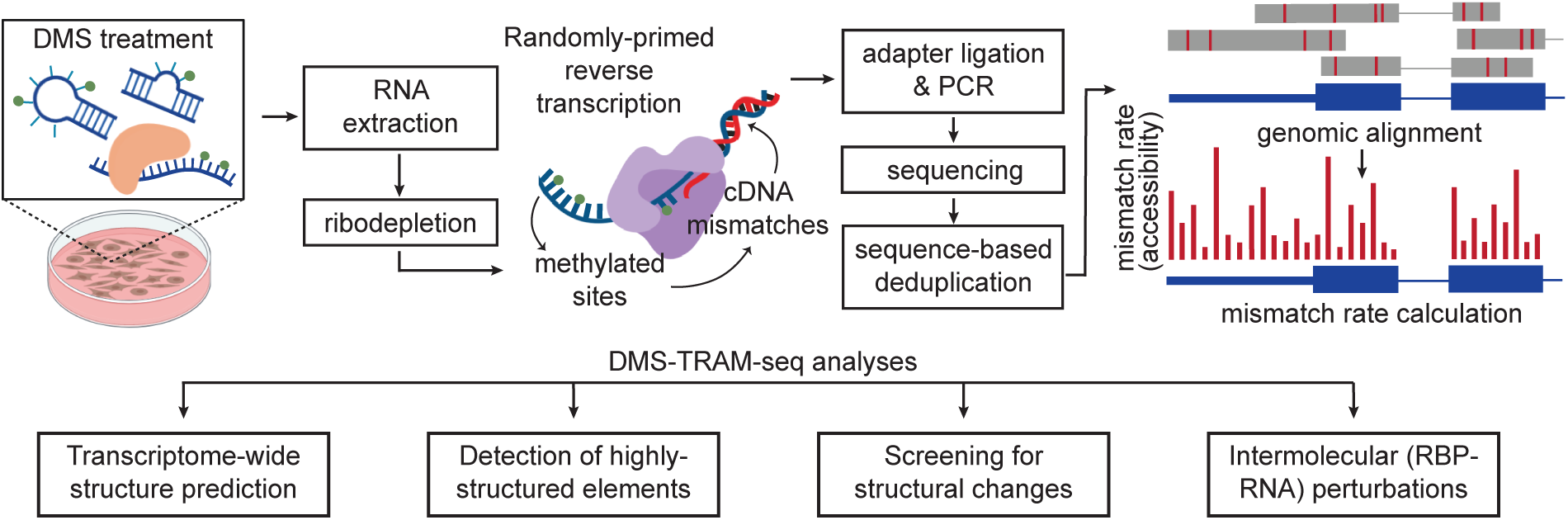
Overview of DMS-TRAM-seq and downstream analyses. DMS modifies RNAs at accessible A and C bases. Extracted RNA, following ribodepletion, is reverse transcribed using group II intron reverse transcriptase (TGIRT). Modified bases produce mismatches during reverse transcription, which are read out using next-generation sequencing. Reads are aligned to the genome to obtain nucleotide-resolution accessibility information. Representative analyses of the mutational profiles using our analysis tools are shown below. DMS-TRAM-seq data provides *in vivo* experimental constraints that can be used for modeling RNA secondary structures at the transcriptome-wide scale. It allows identification of highly structured elements. Comparative analysis across conditions enables the discovery of regions that undergo accessibility changes upon an applied perturbation. Finally, comparison with known RNA-binding protein (RBP) sites facilitates the identification of altered RNA-RBP interactions.

Importantly, the increased coverage of DMS-TRAM-seq allowed us to conduct a quantitative, comparative analysis of RNA accessibilities, enabling us to identify RNAs and their sub-regions that undergo conformational changes in response to an applied perturbation. We demonstrate this capability using oxidative stress as an example, uncovering widespread RNA structural remodeling and perturbed RBP-RNA interactions. Notably, this analysis revealed numerous RBPs that dissociate from their RNA targets during oxidative stress, coinciding with translation inhibition and the relocalization of these RNAs to stress granules. By enabling quantitative, nucleotide-resolution accessibility analysis across the transcriptome, DMS-TRAM-seq uncovers hidden RNA regulatory elements, sheds light on RNA structural dynamics, and provides a powerful platform to examine RNA-mediated gene regulation.

## RESULTS

### DMS-TRAM-seq enables transcriptome-wide RNA structure prediction

Chemical probing approaches, such as DMS-MaPseq, rely on reagents that selectively modify RNA bases depending on their structural context, providing insights into the accessibility and base-pairing status of individual nucleotides. Modified bases produce mismatches during reverse transcription that can be read out via high-throughput sequencing. The resulting mutational profile quantitatively reflects solvent accessibility at single-nucleotide resolution, informing prediction of RNA secondary structure. A high read depth is necessary to accurately ascertain accessibility at a given position on the RNA. As a result, these methods have typically been limited to either the analysis of highly expressed RNAs (such as ribosomal RNAs^31,40^) or use amplicon sequencing that provides high coverage for one or a few regions^37,42^. In order to advance this approach to yield increased coverage across the entire transcriptome, we systematically modified nearly every step in the DMS-MaPseq experimental protocol^35^. Notably, we optimized DMS treatment and RNA extraction conditions, resulting in improved RNA yield and quality (Figure S1A, see Methods). We also modified reverse transcription conditions to facilitate random priming while using the thermostable group II intron derived reverse transcriptase, TGIRT^35^, yielding substantially increased coverage across the transcriptome (Figure 1, S1, see Methods).

The resulting sequencing libraries from DMS-treated and untreated U-2OS cells showed comparable percentages of uniquely mapping reads (79.3% vs. 77.7%, respectively; Figure S1B, see Methods), indicating that DMS treatment does not compromise read quality or mapping efficiency. The mapped reads contained on average 4.88 ± 3.02 (mean ± s.d.) substitutions per 150 × 150 read pair (Figure S1C), corresponding to ∼16.3 modifications per kilobase of RNA. Additionally, the distribution of mismatches across reads was uniform throughout the read length (Figure S1D). The vast majority (86.8%) of mismatches were located at A and C reference bases, as expected by the selectivity of DMS modification (Figure S1E,F). At a relatively stringent threshold of ≥100 quality-filtered mapped reads per base (see Methods), the resulting dataset provided accessibility information for over 21.7 megabases of RNA, representing a significant increase in coverage compared to previous studies. For instance, the original DMS-MaPseq protocol yielded coverage of 733 50-nucleotide regions (36.6 kb total)^35^. Other comparable efforts were similarly limited in scope, such as a SHAPE-MaP study probing Dicer substrates that covered 439 small RNAs^39^ and a recent study probing the 16.5 kb mitochondrial transcriptome^38^.

Our DMS-TRAM-seq data were sufficient for structure prediction of over 9,000 transcripts (Figure 2A,B), representing 71% of the transcriptome expressed in these cells (i.e., genes expressed at ≥ 5 TPM). mRNAs comprised ∼90% of the data (Figure 2B). The dataset also included 265 lncRNAs, despite their generally low expression levels^43^ (Figure 2B, S1G), as well as all 13 mitochondrial protein-coding transcripts and 17 of 22 mitochondrial tRNAs, without requiring mitochondrial isolation as in prior studies^38^ (Table S1). Notably, even without specific optimization for small RNAs, the dataset captured 178 small RNAs (Figure 2B, S1H, Table S1), including 100 snoRNAs (Figure 2B, Table S1).

**Figure 2.**
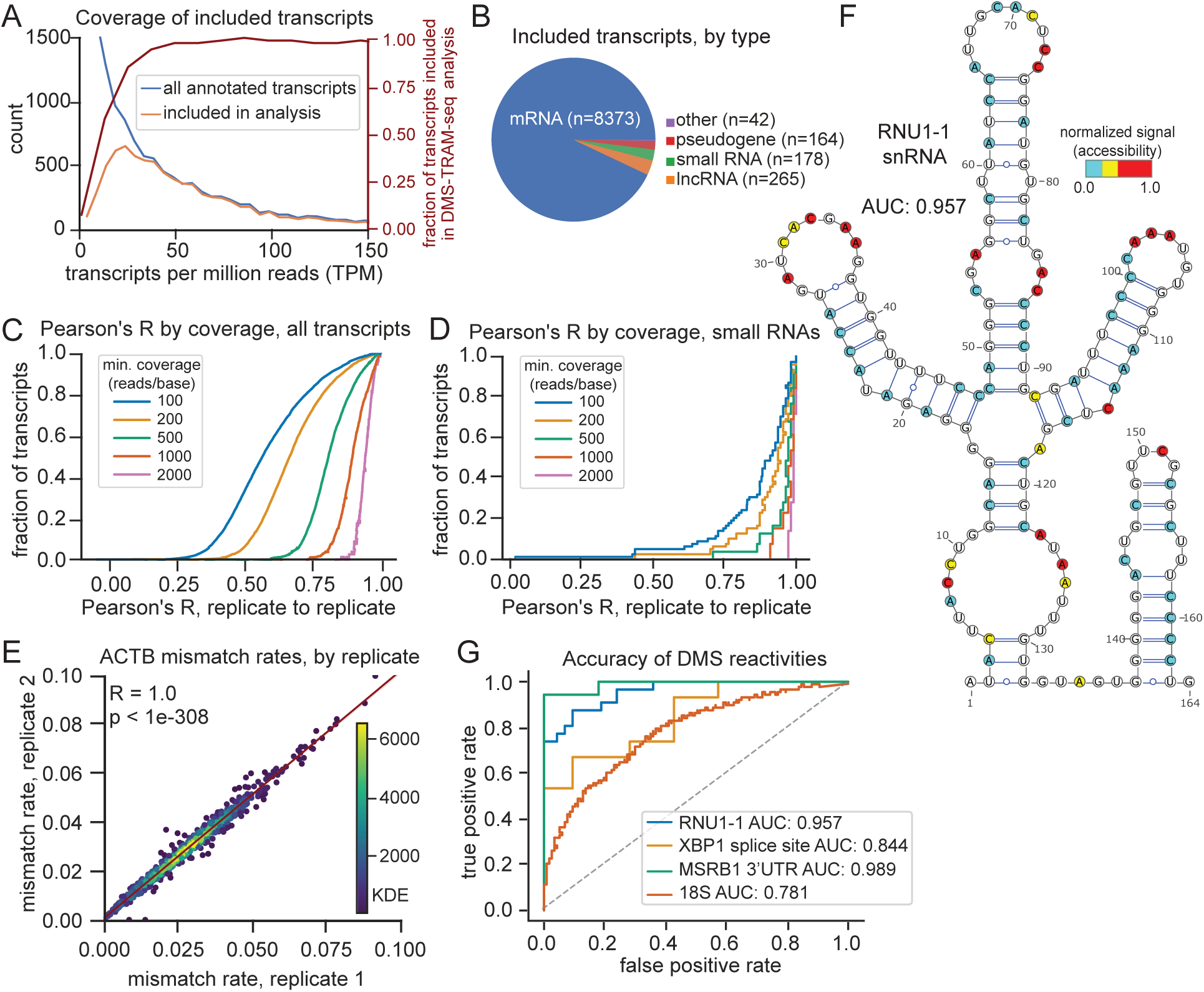
DMS-TRAM-seq provides accurate and reproducible RNA accessibility measurements across thousands of transcripts. (A) Histogram of transcript expression (TPM) for all annotated transcripts in untreated samples (blue) and those with sufficient coverage for DMS-TRAM-seq analysis (orange). The fraction transcripts included in DMS-TRAM-seq analyses at each TPM is shown in red (right y-axis). (B) Distribution of transcript types included in DMS-TRAM-seq analysis, where small RNAs include snoRNAs, scaRNAs, snRNAs, scRNAs, and miRNAs. (C) Distribution of replicate-to-replicate Pearson’s correlation coefficients (R) across all transcripts, where the minimum coverage per nucleotide for a datapoint to be included in the analysis is shown by color. (D) Similar to (C) but only for small RNAs. (E) Scatterplot of mismatch rates across replicates for A/C bases (n = 879) in *ACTB*. Points are colored by density; linear regression shown in red. (F) Overlay of normalized DMS reactivity signal on a published secondary structure model of RNU1-1^37^. (G) Receiver operating characteristic (ROC) curve assessing the alignment of DMS-TRAM-seq signal to known RNA structures. *RNU1-1*, *XBP1* alternate intron, and *MSRB1* 3’UTR element structures are derived from previously-published DMS-MaPseq data^37^, while the 18S rRNA structure is based on PDB 4V6X^90^.

Mutational profiles were highly reproducible between biological replicates, as evidenced by strong correlation between mismatch rates per base across transcripts (Figure 2C, 2D). Correlation was notably higher for RNAs that are expected to be structured (such as snRNAs and snoRNAs) (Figure 2D), consistent with their stable and homogeneous folding (median Pearson’s correlation coefficient, R = 0.892 for small RNAs with coverage ≥ 100 reads/base). mRNAs showed lower overall correlation between replicates (median R = 0.556 at ≥ 100 reads/base), likely reflecting their greater conformational heterogeneity. However, for highly expressed transcripts, such as *ACTB*, we observed highly reproducible signal (R = 1.00, p < 2.2 x 10^-308^, for *ACTB*; median R = 0.805 for RNAs with ≥500 reads/base, Figure 2E), indicating that at sufficient sequencing depth, DMS-TRAM-seq yields robust base-level accessibility profiles and captures the full range of structural conformations, even for RNAs with substantial structural diversity.

We then assessed the accuracy of our mutational profiles by comparing the signal from our dataset to known structures and calculating the area under the receiver operating characteristic curve (AUROC). The AUROC measures how well the DMS-TRAM-seq accessibility signal distinguishes between paired and unpaired bases in a given structure, where an ideal value of 1 indicates perfect alignment and a value of 0.5 indicates random performance. We first examined RNAs that have been previously studied using targeted chemical probing approaches (small nuclear RNA *RNU1-1*^37^, a stem-loop in the 3’UTR of *MSRB1*^44^, and stem-loops in an intron of *XBP1*)^44^. The AUROC for *RNU1-1* was 0.957 (Figure 2F,G, S2A), reflecting high signal accuracy and demonstrating a notable improvement compared to a previous study^37^. The AUROC values for *MSRB1* and *XBP1* were 0.844 and 0.989, respectively, further demonstrating that our accessibility data provides a signal that accurately distinguished paired from unpaired bases (Figure 2G, S2B,C). We also benchmarked our approach against available crystal structures for the *18S* rRNA (PDB: 4V6X) (Figure 2G). The obtained AUROC value of 0.781, while slightly lower than other controls, still displays meaningful alignment with the known crystal structure and is comparable to recent studies^40^. This value likely reflects the intrinsic complexity of rRNA folding and association with ribosomal proteins. Collectively, these validations demonstrate that DMS-TRAM-seq reliably captures base-pairing status and recapitulates known RNA structures across diverse transcripts, underscoring its robustness and utility for transcriptome-wide structural analyses.

Based on our DMS-TRAM-seq data, we provide a publicly accessible database (humanRNAmap.wi.mit.edu) containing predicted secondary structures for 9,022 human transcripts, including 178 small RNAs and 100 snoRNAs, expressed in U-2OS cells. For each transcript, the database includes per-nucleotide DMS reactivity, minimum free energy secondary structure predictions, and base-wise alignment between measured accessibility and predicted base-pairing—enabling users to assess structural confidence and identify well-folded regions. Additional metrics—such as replicate correlation and Gini index—further support the identification of high-confidence structured elements, and are described in detail in later sections. Summary metrics for all transcripts are provided in Table S2.

### DMS-TRAM-seq identifies highly-structured RNAs

To identify RNAs with well-defined secondary structures, we characterized the mutational profile of each transcript using three quantitative metrics. First, we assessed signal reproducibility, as measured by Pearson’s correlation coefficient between replicates. Second, we calculated the Gini index, a commonly used measure of signal uniformity, where a higher value is typically indicative of a structured region^35,41,44,45^. Third, we measured the average mismatch rate as a proxy for the overall accessibility of the region.

Across the transcriptome, increased GC content was associated with modestly lower accessibility and higher Gini values, consistent with the stabilizing effect of GC-rich sequences on secondary structure (Figure S3A). Transcript length, however, did not have any discernable effect on these parameters (Figure S3B). In contrast, various transcript types (such as mRNAs and small RNAs) displayed pronounced differences in the distributions of Gini indices, average accessibilities, and Pearson’s R, with clear clustering among the RNA classes (Figure 3A, S3C). A subset of transcripts exhibited highly reproducible accessibility profiles (Pearson’s R ≥ 0.6) and high Gini indices (≥ 0.4), indicative of well-defined homogeneously folded structures (Figure 3B). These empirically defined thresholds captured multiple known structured RNAs even at their lower limits, including *SNORD3B-1* (R = 0.65, Gini = 0.40) and *SNORA67* (R = 0.63, Gini = 0.46; Figure 3B), validating their use for identifying stably folded transcripts. Applying these criteria, we identified 259 highly-structured transcripts, including 42 snoRNAs, 5 scaRNAs, and two functional ribozymes (Table S3).

**Figure 3.**
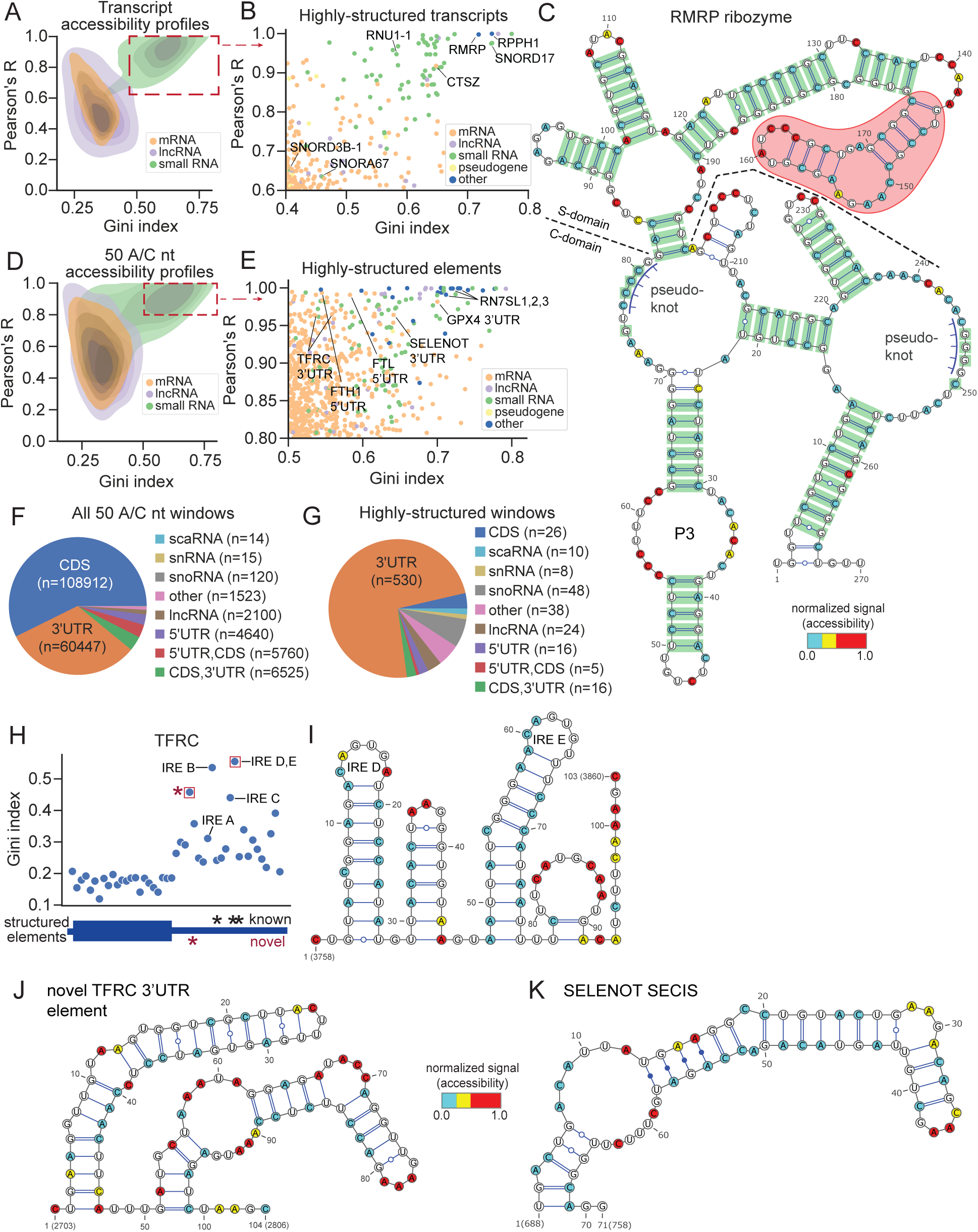
DMS-TRAM-seq identifies structured elements across the human transcriptome. (A) Characterization of transcript accessibility profiles by Gini index and replicate-to-replicate Pearson’s R. n = 9022, see Figure 2 for breakdown by type. (B) Inset of (A) showing the transcripts that are most likely to be structured across their entire length, based on cutoffs of R ≥ 0.6, Gini ≥ 0.4. (C) Predicted secondary structure of the human *RMRP* ribozyme. Base-pairs identified by covariation analysis are in green. The specificity (S) domain is shown above the dashed line; the catalytic (C) domain is below. The structured region shaded in red was not identified by covariation analysis. Pseudoknot prediction via ShapeKnots^91^. (D) Similar to (A) but for 50 A/C nucleotide windows across the transcriptome. n = 190,056. (E) An inset of (C) showing the windows that are most likely to be structured, based on cutoffs of R ≥ 0.8, Gini ≥ 0.5. n = 721. (F) Pie chart showing the breakdown of locations for all analyzed windows in the dataset. n = 190,056. (G) Similar to (F), but only for the highly-structured windows shown in (C). (H) Gini indices for 50 A/C nucleotide windows tiled across the canonical TFRC transcript. Known iron response elements (IREs) and a newly identified structured region are annotated below. Red boxes indicate windows analyzed in (I) and (J). (I) Structure prediction for the window with the highest Gini index in the TFRC 3’ UTR, containing two known IREs (IRE-D and IRE-E). (J) Predicted structure of a novel TFRC 3′UTR element not overlapping any annotated IRE. (K) Predicted secondary structure of the selenocysteine insertion element (SECIS) in SELENOT. Noncanonical A–G base pairs were manually incorporated into the RNAstructure Fold output.

One of the most highly-structured transcripts that emerged from our analysis is the RNA component of mitochondrial RNA processing endoribonuclease (*RMRP*) ribozyme (Figure 3B,C). This non-coding RNA is evolutionarily conserved across eukaryotes^46–48^ and plays a critical role in rRNA processing by cleaving the intervening sequence (ITS1) between the *18S* and *5.8S* rRNAs^49,50^. The RMRP ribozyme contains a catalytic and a specificity domain^51^, and mutations in this gene are associated with numerous genetic disorders^52,53^. Despite its functional importance, structural information on human *RMRP* has remained limited, aside from one highly-conserved stem-loop (P3 domain)^54^, though a recent study provided the cryo-EM structure of the *S. cerevisiae RMRP* homologue^51^. We used our DMS-TRAM-seq data to generate a secondary structure model of human *RMRP* (Figure 3C). Notably, the P3 domain in our predicted structure exhibits a perfect alignment with the known crystal structure^54^. Furthermore, we found extensive secondary structure similarity between the catalytic domains of yeast and human *RMRP*, despite only 52% sequence similarity (Figure S4A).

To further validate our structural model, we performed evolutionary covariation analysis across 933 eukaryotic *RMRP* homologs^55,56^ (Figure S4B). This analysis identified 63 significantly covarying base-pairs^57^ (E < 0.05, see Methods), of which 62 were predicted to form base-pairs in our DMS-TRAM-seq derived model (Figure 3C, highlighted in green). The remaining covarying pair (G-75 to C-250), while available for pairing adjacent to a predicted pseudoknot, was not explicitly predicted in our structural model.

Covariation analysis, while powerful for predicting RNA secondary structures, is inherently limited by its reliance on identifying evolutionary covariation within homologous regions across multiple species. In our structural model of *RMRP*, the covarying pairs accounted for 79% of all predicted base pairs. However, DMS-TRAM-seq data identified several base-pairs not detectable by covariation analysis, including an additional stem-loop structure within the specificity domain (Figure 3C, indicated in red). This previously uncharacterized element likely contributes to target recognition by human *RMRP*. These results showcase the power of DMS-TRAM-seq in capturing high-confidence RNA structures at a transcriptome-wide scale. By directly probing RNA base accessibility in living cells, DMS-TRAM-seq complements covariation analysis and high-resolution structure determination methods, yielding a comprehensive view of RNA folding in its native cellular context.

### Identification of highly-structured RNA elements

Most transcripts, particularly mRNAs, are expected to contain relatively small regions that fold into defined structured elements. To identify such locally structured regions, we split each transcript into small, non-overlapping windows of 50 quality- and coverage-filtered A/C bases, yielding a total window size of 103 ± 25 nucleotides (mean ± s.d., see Methods, Figure S5). Our DMS-TRAM-seq dataset covered ∼190,000 windows, spanning ∼20 megabases of RNA (Figure 3D, Table S4). In order to identify windows that are highly likely to harbor structured elements, we calculated the replicate-to-replicate Pearson’s correlation coefficient (R) for each window and selected those exhibiting low variability (R ≥ 0.8) along with a high Gini index (≥ 0.5) (Figure 3E). These thresholds are more stringent compared to those used in the analysis of full-length transcripts and captured known structured elements even at the lower end of these ranges, such as stem-loop structures within *SNORD3D* (Gini index = 0.537, R = 0.807) and *SNORD84* (Gini index = 0.492, R = 0.791).

Using these cut-offs, we identified 721 regions across the transcriptome that are likely to harbor structured elements (Table S5), representing the top 0.38% of all windows. The majority (593) were located within mRNAs, with a notable enrichment in 3’UTRs: 546 of the structured elements (92.4%) mapped to or partially overlapped with 3’UTRs, despite these regions constituting only 35% of the dataset (p = 1.97 × 10^-109^, hypergeometric test) (Figure 3F,G). In contrast, structured elements were significantly underrepresented in CDSs (47 windows, 6.5%; p = 1.09 × 10^-233^) and 5′UTRs (21 windows, 2.9%; p = 3.69 ×10⁻⁴) (Figure 3F,G), suggesting that RNA folding is disfavored in these regions. The observed enrichment of structured elements in 3’UTRs aligns with their established role as hubs for post-transcriptional regulatory interactions, often mediated by RNA structure^58^.

As an example, we analyzed the *TFRC* transcript, which contained multiple regions that met our stringent criteria for identifying highly structured elements. *TFRC* encodes for the transferrin receptor, a key mediator of cellular iron uptake. Its expression is tightly regulated by iron response elements (IREs) in its 3’UTR, which bind iron regulatory proteins (IRPs) to modulate mRNA stability and translation in response to cellular iron levels^59,60^. Gini index analysis across the *TFRC* transcript revealed at least four windows with substantial secondary structure within the 3′UTR, in contrast to the relatively unstructured CDS and 5’UTR (Figure 3H). We used the accessibility constraints from our DMS-TRAM-seq data to model the secondary structure for the window with the highest Gini index and observed two known IREs within this region. Our predicted folds for these IREs precisely matched their established structures (Figure 3I)^60^. All other known IREs in *TFRC* were also observed in high-Gini windows. In addition to these known elements, we also identified a previously uncharacterized structured element that does not overlap with any annotated IRE (Figure 3H,J), suggesting the presence of another putative regulatory motif. Although 5′UTRs are generally depleted of stable secondary structure, our analysis also captured two well-characterized IREs in the 5′UTRs of *FTH1*^61^ and *FTL*^62^, both of which are known to adopt defined RNA folds.

Selenoprotein transcripts provide another example of RNAs that rely on highly conserved structural elements. These mRNAs encode a unique class of proteins that incorporate selenocysteine (Sec), the 21st amino acid, essential for redox homeostasis and metabolic regulation^63^. Unlike standard amino acids, Sec is co-translationally inserted at UGA codons, which normally signal translation termination^63^. This recoding is directed by a selenocysteine insertion sequence (SECIS), a highly conserved stem-loop structure located in the 3’UTR of selenoprotein mRNAs, which helps recruit a charged Sec-tRNA to the elongating polypeptide, ensuring selenocysteine incorporation instead of termination^64^.

Consistent with this intricate structural requirement, we identified several highly structured windows in selenoprotein transcripts (Figure 3E). For example, we detected a prominent structured element in the 3’UTR of *SELENOT*, consistent with the presence of a SECIS motif (Figure 3E,K). SECIS elements are characterized by distinct structural features, including tandem noncanonical adenine-guanine (A-G) pairs^65^ that form kink-turn motif, and conserved apical loop adenines, which serve as key recognition sites for SECIS-binding protein 2 (SBP2)^64^. These noncanonical interactions are rarely captured by sequence-based structure prediction algorithms, which generally account only for Watson–Crick and G–U base pairs^66^. While DMS-TRAM-seq does not explicitly predict noncanonical base pairing, the underlying accessibility profiles accurately reflect reduced reactivity in these regions, enabling detection of stable structures that elude standard algorithms (Figure 3K).

Besides these specific examples, the expanded coverage enabled by DMS-TRAM-seq revealed hundreds of regions likely to adopt defined secondary structures in cells. Using our stringent thresholds (R ≥ 0.8, Gini ≥ 0.5), we identified 721 high-confidence structured elements across 602 transcripts, spanning diverse RNA classes (Table S5). Additional structured regions likely exist within the dataset, and their secondary structures can be modeled using DMS-TRAM-seq’s experimentally derived accessibility constraints. Accessibility profiles across the entire transcriptome, segmented into defined windows, and their secondary structure predictions are available through our interactive web portal.

### Translation inhibition triggers global RNA folding

Previous studies have shown that translation can remodel RNA structure, and actively translated mRNAs are generally unfolded in the cell^8,44,67^. The increased coverage of DMS-TRAM-seq allowed us to systematically investigate these effects. Consistent with ribosomal scanning during translation initiation and elongation, we observed that 5’UTRs and CDSs are significantly more accessible than 3’UTRs (Figure 4A). To further explore this relationship, we integrated DMS-TRAM-seq data with a ribosome profiling dataset from the same cell type^68^ and uncovered a clear relationship between translation efficiency (TE) and RNA accessibility. Transcripts with high ribosome occupancy (TE ≥ 2) exhibited markedly higher accessibility in their 5’UTRs and CDSs compared to those with low ribosome occupancy (TE ≤ 0.5) (Figure 4B). In contrast, accessibility within the 3’UTRs was comparable across the TE groups (Figure 4B), consistent with translation-dependent remodeling being confined to the regions directly engaged by the ribosome.

**Figure 4.**
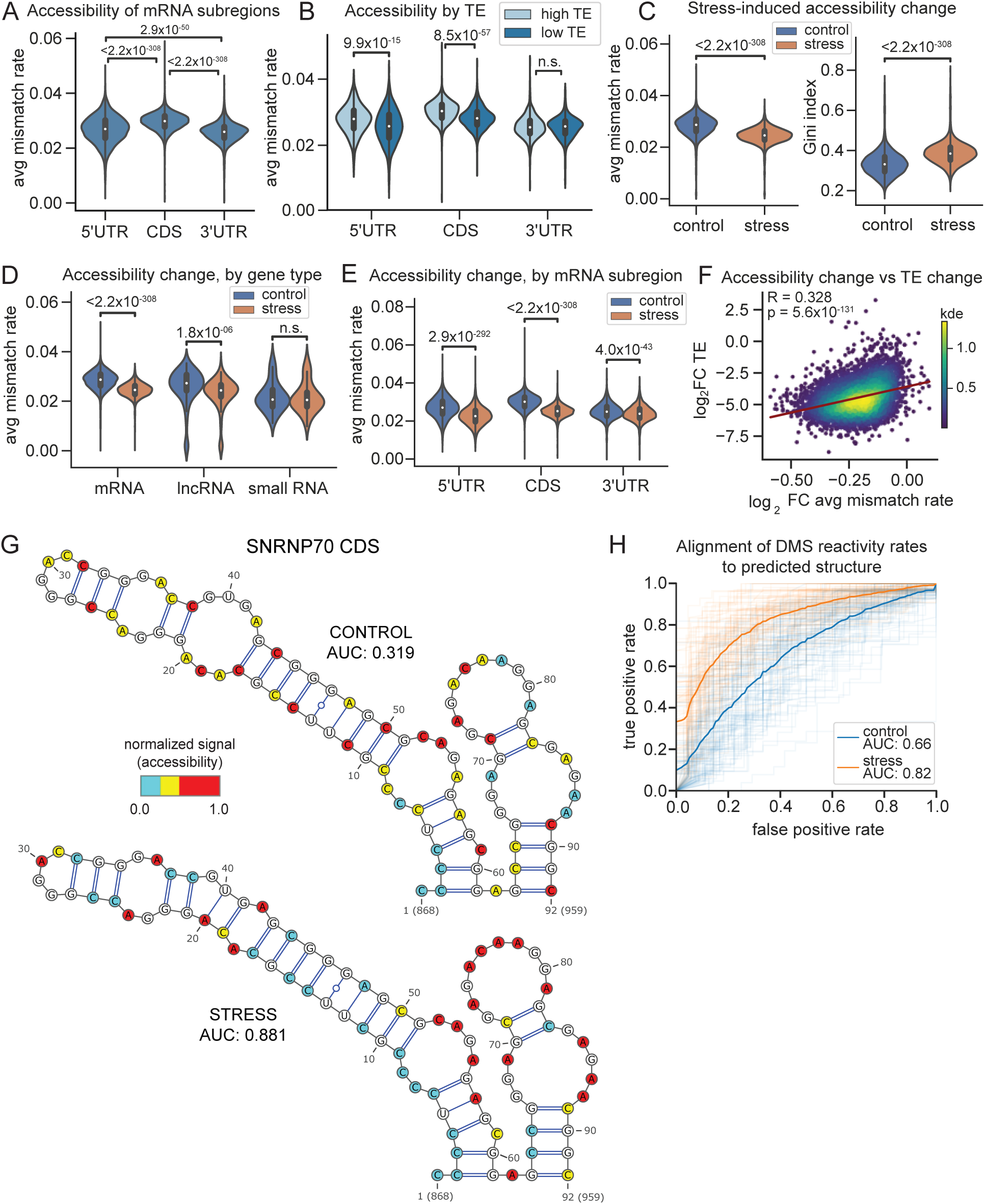
RNAs undergo increased folding into thermodynamically favored conformations upon translation inhibition. (A) Distribution of average mismatch rates across different mRNA subregions. (B) Same as (A), but where transcripts are stratified by their translation efficiencies (TE) from ribosome profiling data^68^. High TE transcripts have a TE ≥ 2, while low TE transcripts have a TE ≤ 0.5. (C) Distributions of average mismatch rates (left) and Gini indices (right) calculated for each transcript, in control (blue) and oxidative stress (orange) conditions. (D) Distributions of average mismatch rates across different transcript types, under control (blue) and oxidative stress (orange) conditions. Small RNAs include snoRNAs, scaRNAs, snRNAs, scRNAs, and miRNAs. (E) Similar to (D) but for different mRNA subregions. (F) Correlation between transcript-wide changes in RNA accessibility (average mismatch rate across a transcript) and TE after oxidative stress^74^. Datapoints colored by density. (G) Structure prediction for a region in the *SNRNP70* CDS overlaid with DMS signal. AUC quantifies agreement between the DMS signal and the predicted structure. (H) ROC curves comparing DMS signal to the predicted structure for the top 100 regions undergoing structural changes after oxidative stress. Thin lines represent individual regions in control (blue) and oxidative stress (orange) conditions, respectively. Bold lines indicate the average ROC curve for each condition, with corresponding AUROC values.

We next examined how RNA accessibility changes upon translation inhibition using sodium arsenite, a potent oxidative stress inducer. Sodium arsenite rapidly inhibits translation initiation via phosphorylation of eIF2α^69^, allowing for the observation of immediate effects on RNA structure without directly interfering with translation elongation or artificially stalling ribosomes on mRNAs as with other translation inhibitors. Arsenite treatment also induces the formation of stress granules — membraneless ribonucleoprotein assemblies enriched in untranslated mRNAs and RNA-binding proteins (RBPs)^70–72^. Notably, previous studies have shown that stress granule formation relies on extensive RNA-RNA interactions^73^. Given DMS-TRAM-seq’s ability to provide transcriptome-wide coverage with single nucleotide resolution, we reasoned that it could serve as a powerful tool to examine these structural rearrangements at scale.

As expected, treatment with 0.5 mM sodium arsenite for 30 minutes resulted in the accumulation of stress granules, as observed by relocalization of the marker protein G3BP1 (Figure S6A,B). Importantly, DMS treatment in arsenite-treated cells had no discernible effect on stress granule morphology (Figure S6C,D), indicating that chemical probing does not perturb granule integrity. DMS-TRAM-seq in untreated and arsenite-treated U-2OS cells yielded robust coverage for 6,396 transcripts across both conditions (see Methods). Strikingly, oxidative stress led to a marked transcriptome-wide decrease in RNA accessibility, reflected by reduced mismatch rates and a concurrent increase in Gini indices, indicating a global transition to more structured RNA conformations (Figure 4C). These effects were most pronounced in mRNAs, followed by lncRNAs, while small RNAs remained largely unaffected (Figure 4D, S7A). Within mRNAs, accessibility decreased most in 5′UTRs and CDSs, with relatively modest changes in 3’UTRs (Figure 4E, S7B), consistent with reduced ribosome engagement in regions normally involved in initiation and elongation.

To further investigate how translation impacts RNA structure under stress, we integrated our DMS-TRAM-seq data with ribosome profiling data from arsenite-treated U-2OS cells^74^. For each transcript, we compared the change in RNA accessibility with the change in translation efficiency following arsenite exposure. This analysis revealed a significant correlation between reduced translation and increased RNA folding: transcripts with the largest decreases in TE also showed the greatest reductions in accessibility (R = 0.328, p = 5.6 × 10⁻¹³¹) (Figure 4F). Altogether, these results highlight widespread RNA structural remodeling in response to oxidative stress and support the view that active translation suppresses RNA folding in cells.

We next examined whether these structural changes were linked to stress granule localization. Using a published dataset of stress granule-enriched RNAs^72^, we found that transcripts excluded from stress granules exhibited the largest reductions in accessibility upon arsenite treatment (Figure S7C). This trend mirrors their strong translational repression under oxidative stress^74^. Stress granule-enriched transcripts also showed reduced accessibility upon arsenite treatment, though this change was less substantial (Figure S7C). Interestingly, these stress granule-enriched transcripts displayed lower baseline accessibility prior to stress exposure (Figure S7D), suggesting that stress granule-associated transcripts tend to be more structured even in unstressed conditions. Together, these observations indicate that RNA folding following oxidative stress is primarily driven by translation inhibition, rather than by stress granule formation itself. mRNAs excluded from granules may undergo increased folding due to the loss of ribosome-mediated unwinding and the displacement of translation-associated RBPs such as eIF4A^75^. In the absence of these proteins, previously constrained regions may adopt thermodynamically favored conformations. In contrast, granule-enriched transcripts are already more structured prior to stress exposure, suggesting that pre-existing RNA structure may contribute to their selective recruitment into stress granules. These RNAs undergo relatively modest structural remodeling upon stress, consistent with a model in which granule association either preserves existing structures or protects RNAs from further folding.

### RNAs fold into their thermodynamically favored structures upon translation inhibition

DMS-TRAM-seq also enabled transcript-level analysis of RNA structure remodeling under stress. For instance, *ACTB*, one of the most abundant mRNAs in the cell, adopts a more structured conformation under arsenite-induced oxidative stress, as evidenced by a marked reduction in accessibility (Figure S7E). We modeled the secondary structure of *ACTB* RNA utilizing DMS-TRAM-seq data as constraints and found that, while the predicted structures before and after stress remained largely similar, their agreement with the underlying DMS signal improved in the stress condition (Figure S7F). Strikingly, under stress conditions, the predicted structure was identical to that predicted solely from sequence, without incorporating DMS constraints (Figure S7F). This convergence suggests that inhibition of translation initiation allows *ACTB* to fold into its thermodynamically favored structure, likely unmasked by the absence of ribosome-mediated remodeling.

Given this striking effect on *ACTB*, we next asked whether similar structural shifts occur more broadly across the transcriptome. To address this question, we examined transcripts that exhibited the most pronounced changes in accessibility upon arsenite treatment (see below). One of the strongest shifts was observed within the coding sequence of SNRNP70 (Figure 4G), a highly-expressed protein subunit of the U1 snRNP complex. Despite minimal differences in predicted secondary structures between control and stress conditions, their agreement with the underlying DMS signal improved substantially. In untreated cells, highly reactive bases were often predicted to be paired, yielding a high “false positive” rate and therefore a low AUROC value (Figure 4G, top, AUROC = 0.319). Under arsenite stress, however, the agreement between structural prediction and underlying DMS-TRAM-seq data improved dramatically, with low-reactivity bases consistently mapping to paired regions and vice versa, yielding a much higher AUROC value of 0.881 (Figure 4G, bottom).

We expanded this analysis to the top 100 transcript regions with the largest stress-induced changes in accessibility (see Methods). In nearly all cases, the agreement between predicted secondary structure and DMS-TRAM-seq signal improved following stress (Figure 4H). These results suggest that RNA folding becomes more consistent with thermodynamic predictions upon translation inhibition, supporting a model in which ribosome disengagement allows transcripts to adopt their native conformations. Together, these results underscore the tight coupling between translation and RNA structural rearrangements under oxidative stress, and highlight DMS-TRAM-seq’s ability to capture these dynamic changes at an unprecedented scale and resolution.

### Systematic identification of RNA regions undergoing structural remodeling

We hypothesized that the high coverage provided by DMS-TRAM-seq could allow us to systematically identify RNAs that undergo conformational changes in response to an applied perturbation. To facilitate this analysis, we segmented the transcriptome into non-overlapping windows, each containing 50 high-confidence A/C bases (as described above; see Methods). For each window, we computed a delta (δ) score, defined as the average bidirectional change in per-nucleotide reactivity between conditions (Figure 5A). This metric captures both increases and decreases in reactivity, enabling detection of local structural remodeling, in addition to general increases or decreases in folding.

**Figure 5.**
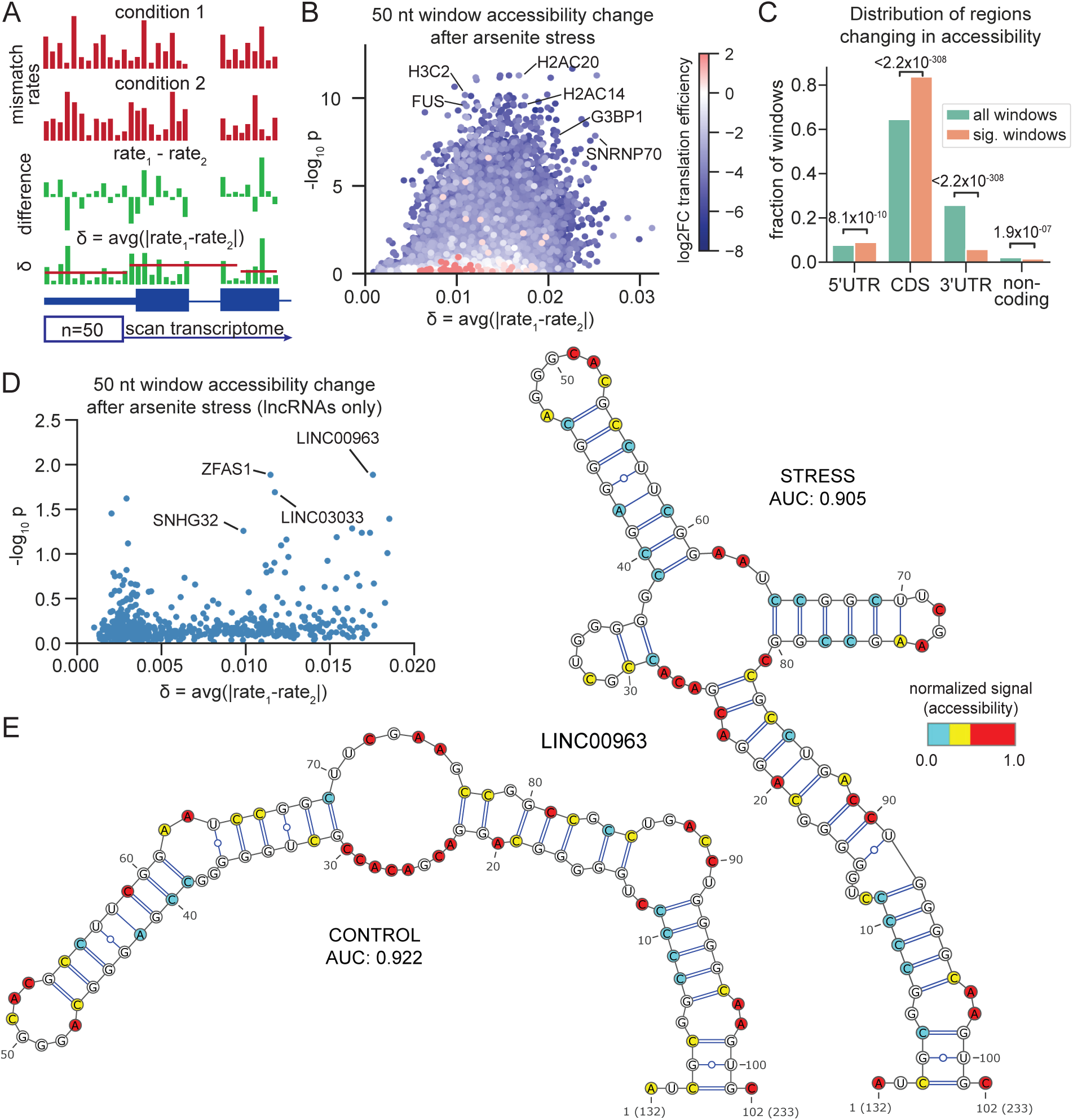
Comparative analysis of mutational profiles identifies RNA regions undergoing significant accessibility changes under stress. (A) Schematic of transcriptome-wide screening approach, where 50 A/C nucleotide windows are analyzed to quantify both the magnitude (δ) and the significance (p-value) of change in accessibility between control and treatment conditions. δ is the bidirectional change in mismatch rate per base, averaged over the entire window. (B) Identified regions based on the analysis in (A) displaying significant structural change under oxidative stress. Points are colored by change in TE after oxidative stress^74^. (C) Distribution of windows that are significantly changing in accessibility (orange) after arsenite stress, compared to all analyzed windows (teal). P-values are calculated using hypergeometric test. (D) Similar to (B), but limited to windows within lncRNAs. (E) Secondary structure prediction for the *LINC00963* window identified in (D), comparing control (left) and the arsenite stress (right) conditions.

To assess statistical significance, we compared the within-condition reproducibility of DMS reactivity profiles (Pearson correlation between replicates) to the between-condition correlation coefficients, using a two-sample *t*-test. This approach detects regions where mutational profiles are stable within each state but differ substantially between conditions, indicating reproducible and condition-specific structural rearrangements. Windows with a high δ and low p-value reflect regions that undergo substantial conformational rearrangements with high confidence, based on both magnitude and reproducibility of change.

We applied this analysis to identify the windows that exhibit statistically-significant (p_adj_ ≤ 0.05) changes in accessibility upon arsenite treatment (Figure 5B; Table S6). Windows showing significant structural changes were enriched in coding sequences and 5’UTRs, while being relatively depleted in 3’UTRs (Figure 5C). This trend aligns with our earlier observation that translation inhibition is a major driver of RNA structural rearrangements after arsenite stress. Notably, we identified regions undergoing substantial accessibility changes in the coding sequences for *FUS*, *G3BP1*, *SNRNP70*, and several histone mRNAs (Figure 5B), all of which exhibit pronounced decreases in translation efficiency under oxidative stress^74^.

Since translation dynamics play a dominant role in shaping mRNA structures, we next asked whether DMS-TRAM-seq could also detect subtler rearrangements in noncoding RNAs, which are not subject to the same ribosome-associated effects. As a proof of concept, we analyzed the lncRNA *NORAD*, which relocalizes to stress granules upon arsenite treatment^72^. A recent study revealed that this relocalization is accompanied by structural changes at its 3’ end^42^. Our DMS-TRAM-seq data recapitulated these structural rearrangements, revealing similar changes in RNA accessibility under stress (Figure S7G).

We extended this approach to systematically evaluate structural remodeling across lncRNAs. While changes in accessibility were generally more modest than those observed in protein-coding transcripts, several lncRNA regions exhibited significant stress-induced rearrangements (Figure 5D). One notable example is *LINC00963*, a lncRNA that partially localizes to stress granules^72^. Structure prediction using DMS-TRAM-seq constraints revealed substantial remodeling between control and stress conditions, with more than half of the predicted base pairs (17 of 33) differing between states (Figure 5E). The accessibility data aligned closely with predicted structures in both conditions (AUROC = 0.922 in control; 0.905 under stress), suggesting that *LINC00963* adopts distinct, well-defined conformations under oxidative stress. While the functional consequence of this rearrangement is unclear, such transitions may influence—or result from—changes in RBP binding and stress granule recruitment. More broadly, these results highlight the ability of DMS-TRAM-seq to uncover condition-specific RNA structures at single-nucleotide resolution. This comparative analysis approach provides a powerful new framework to identify RNAs undergoing conformational remodeling in response to a given cellular perturbation.

### Identifying perturbed RBP-RNA interactions under oxidative stress

We next examined whether changes in RNA-binding protein (RBP) association could be inferred from DMS-TRAM-seq data. One of the key proteins involved in the cellular response to arsenite-induced oxidative stress is G3BP1, a single-stranded RNA-binding protein that, along with its homologue G3BP2, is essential for stress granule formation^76^. A recent study showed that G3BP1 dissociates from a majority (>75%) of its target transcripts upon exposure to oxidative stress^77^, offering a benchmark for assessing whether DMS-TRAM-seq captures RBP-RNA dynamics at nucleotide resolution.

To capture localized changes associated with RBP binding, we examined the accessibility profiles of 20 A/C nucleotide windows (as described above) in order to account for the typical footprint of RBPs^78^ (Table S7). We observed a modest but statistically significant shift in accessibility at G3BP1 dissociation sites relative to retained sites (Figure S8A,B). Since accessibility changes in coding sequences and 5’UTRs may be influenced by translation inhibition, we focused our analysis on G3BP1 binding sites within 3’UTRs. Strikingly, sites where G3BP1 dissociates showed pronounced changes in accessibility (as measured by our δ metric) compared to sites where G3BP1 remains bound (Figure 6A). Notably, dissociated sites had significantly lower accessibility after oxidative stress, reflecting increased RNA folding after G3BP1 release, while bound sites showed no significant change (Figure 6B). This result supports a model in which G3BP1 binding prevents local RNA folding; once G3BP1 dissociates under stress, the freed RNAs are able to adopt a more folded conformation, likely participating in intra- and inter-molecular RNA interactions that may foster stress granule formation^79,80^.

**Figure 6.**
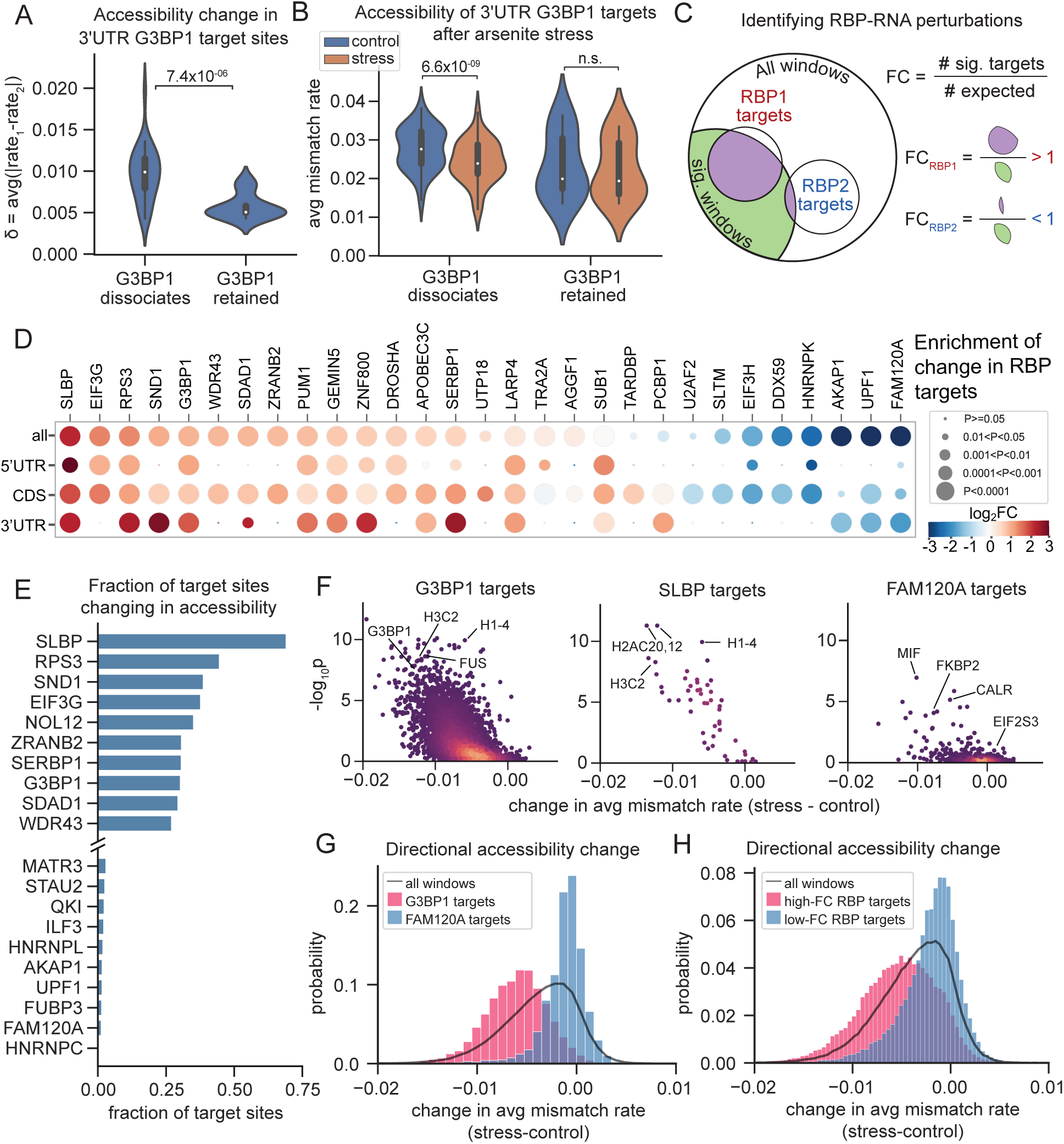
DMS-TRAM-seq identifies altered RBP–RNA interactions under oxidative stress. (A) Change in mismatch rate (δ) for G3BP1 target sites in 3’ UTRs, where G3BP1 either dissociates (left) or remains bound (right) after oxidative stress^77^. (B) Average mismatch rates for the same regions as in (A) under control (blue) and oxidative stress (orange) conditions. (C) Schematic illustrating the analysis used to identify RBPs whose target sites contain a disproportionately large (“RBP1”, red) or small (“RBP2”, blue) fraction of target sites significantly changing in DMS signal. (D) Heatmap showing enrichment (red) or depletion (blue) of significant structure changes within the target sites of the indicated RBPs, where the fold change value is calculated as indicated in (C), relative to a randomized null model (see Methods). P-values calculated by the hypergeometric test. (E) Fraction of target sites for each RBP that are significantly changing in structure; only the top 10 and bottom 10 RBPs are shown. (F) Directional changes in accessibility for target sites of G3BP1 (left), SLBP (middle), and FAM120A (right), where negative values indicate reduced accessibility in the stress condition. Points are colored by density. (G) Distribution of directional accessibility changes for target sites of G3BP1 (red), FAM120A (blue), and all windows (black line). (H) Similar to (G), but comparing all windows (black) to windows overlapping target sites of RBPs with high (red) and low (blue) fold-change scores from (D).

To systematically identify RBPs whose RNA-binding activity is altered during oxidative stress, we integrated our DMS-TRAM-seq data with CLIP (cross-linking and immunoprecipitation) datasets for ∼150 RBPs from ENCODE^81^. For each RBP, we assessed whether its target sites were overrepresented among transcript regions with significant accessibility changes following arsenite treatment. Fold enrichment was calculated relative to a shuffled null model (Figure 6C; see Methods), and significance was assessed using the hypergeometric test. To account for potential differences in untranslated regions versus coding sequences, we performed these analyses for target sites across the full transcript, as well as within specific regions (5′UTR, CDS, and 3′UTR). RBPs whose target sites were most enriched or depleted in accessibility changes (|log_2_FC| ≥ 1.1, p_adj_ ≤ 0.001) are shown in Figure 6D (for full results, see Table S8).

Among the RBPs with the strongest shifts in accessibility at their target sites were SLBP, eIF3G, and RPS3. SLBP binds a conserved stem-loop at the 3′ end of replication-dependent histone mRNAs, which lack poly(A) tails. Following oxidative stress, ∼80% of SLBP-associated sites exhibited significant accessibility changes (Figure 6E), far exceeding the transcriptome-wide background rate of 11.3%. As with G3BP1 targets, the vast majority of SLBP-associated sites exhibited reduced accessibility (Figure 6F-H), consistent with an increase in RNA secondary structure. These findings suggest that SLBP, like G3BP1, may dissociate from its targets under stress, potentially altering histone mRNA stability or processing.

eIF3G is a core subunit of the eukaryotic initiation factor 3 (eIF3) complex, and is essential for recruiting the 40S ribosomal subunit to mRNA. Previous studies have shown that eIF3G dissociates from its target mRNAs under stress, disrupting the scanning process and inhibiting translation initiation^82^. Similarly, RPS3, a ribosomal protein located near the mRNA entry site on the 40S ribosomal subunit, stabilizes mRNA-ribosome interactions during translation initiation and elongation. The observed accessibility changes at eIF3G and RPS3 binding sites are consistent with global translational suppression under oxidative stress^82^ and reflect a transition toward increased secondary structure as these RBPs dissociate from their targets.

In contrast, several RBPs showed significantly fewer accessibility changes at their target sites than expected (Figure 6D, G-H, in blue). These included HNRNPK, UPF1, and FAM120A, all of which are known to relocalize to stress granules^76,83–86^. The relative stability of their binding sites suggests that these RBPs may remain bound to their RNA targets during oxidative stress, preventing structural remodeling that would otherwise occur. Alternatively, they may preferentially bind to pre-folded or structurally stable RNAs that are less susceptible to stress-induced refolding. In either case, DMS-TRAM-seq captures the differential behavior of RBPs during stress, revealing both sites of dissociation and regions of retained interaction.

Taken together, these results demonstrate that DMS-TRAM-seq can sensitively capture changes in RNA accessibility associated with RBP dissociation or retention, offering a transcriptome-wide view of dynamic RNA–protein interactions. By integrating accessibility profiles with binding site data, DMS-TRAM-seq provides a powerful framework to interrogate how cellular perturbations affect the complex network of interactions that shape RNA structure and function.

## DISCUSSION

DMS-TRAM-seq enables true transcriptome-wide RNA accessibility profiling, allowing for high-resolution mapping of the RNA structurome. Our optimized methods captured the solvent accessibility information at nucleotide resolution for the majority of expressed transcripts, with substantial gains in both coverage and data quality over prior approaches. Our dataset provided sufficient coverage for accessibility analysis on >70% of all expressed transcripts, and over 95% of those expressed above 25 TPM, without the need for sample pooling or data extrapolation. This accessibility information provides experimentally determined constraints for secondary structure prediction. Our analysis also revealed >700 previously unannotated, high-confidence structured elements, many of which may serve regulatory roles. While structured elements such as riboswitches are well characterized in bacteria, their mammalian counterparts remain elusive^87^. DMS-TRAM-seq allows for the systematic discovery of such structured elements in human transcripts. To support community use, we provide an open-access database covering ∼20 megabases of RNA sequence, allowing users to generate *in vivo*-constrained secondary structure models for any region of interest.

Covariation analysis remains the gold standard for identifying conserved RNA structures but is inherently limited to regions with sufficient evolutionary depth and coevolving base pairs. Many biologically important RNA elements lack the number or diversity of homologs needed for robust covariation analysis. DMS-TRAM-seq overcomes these limitations by providing direct, *in vivo* measurements of RNA accessibility, enabling structural analysis even in transcripts that are poorly conserved, sparsely annotated, or do not contain sufficient covarying pairs for structure prediction. As demonstrated with human RMRP, integrating DMS-TRAM-seq with covariation analysis yields more complete structural models. Beyond comparative approaches, the quantitative profiles from DMS-TRAM-seq can serve as structural constraints or training input for emerging deep learning models. While tools like AlphaFold^88^ have transformed protein structure prediction, progress in RNA modeling has been limited by the scarcity of experimental datasets. The experimentally constrained secondary structures from DMS-TRAM-seq may serve as valuable inputs for deep learning approaches, advancing the modeling of RNA tertiary structures with greater accuracy.

The nucleotide-resolution accessibility information from DMS-TRAM-seq can also be integrated with other high-throughput datasets to uncover structure-function relationships. By combining DMS-TRAM-seq data with ribosome profiling, we uncovered that translation actively remodels mRNA structures and, upon translation inhibition, mRNAs tend to fold into their native, thermodynamically favored conformations. Integration with CLIP-seq data allowed us to discover perturbed RBP-RNA interactions following cellular stress. These analyses provide a proof of principle, demonstrating that DMS-TRAM-seq can be integrated with diverse datasets, including RBP motifs, miRNA target sites, and RNA modification maps, to uncover how distinct layers of post-transcriptional regulation relate to RNA conformational changes in cells.

The high coverage provided by DMS-TRAM-seq enables comparative, transcriptome-wide analysis of RNA accessibility across conditions, allowing us to nominate transcripts that undergo conformational changes in response to cellular perturbations. These shifts may reflect altered RNA folding, protein binding, or localization, offering insights into how RNAs respond to regulatory cues. This ability to resolve transcript-specific conformational dynamics opens new avenues for investigating post-transcriptional regulation in both normal and diseased states. For example, DMS-TRAM-seq can be applied in disease models to detect transcripts affected by mislocalized or mutant RBPs. This is particularly relevant in neurodegenerative disorders such as ALS and FTD, where mutations in proteins like TDP-43 and FUS are linked to widespread RNA misprocessing. By profiling accessibility changes, DMS-TRAM-seq can uncover candidate RNAs whose regulation is disrupted in disease, providing new insight into RNA-based mechanisms of pathogenesis and highlighting potential therapeutic targets.

While DMS-TRAM-seq enables transcriptome-wide secondary structure prediction and the identification of highly structured elements, it captures an ensemble average of RNA conformations within the cell. RNAs may adopt multiple conformations that may be essential for their regulatory functions, such as in the case of riboswitches^89^. Likewise, rapid structural transitions due to translation or changes in RBP binding may not be captured in our current iteration of DMS-TRAM-seq. Combining DMS-TRAM-seq with long read sequencing technologies and clustering-based analyses^37^ may help tease apart RNAs that exist in multiple coexisting conformations. As a DMS-based approach, DMS-TRAM-seq primarily probes single-stranded A and C residues. Additionally, RNA-binding proteins (RBPs) or extensive tertiary interactions can shield regions from modification by DMS, complicating interpretation.

Despite these considerations, DMS-TRAM-seq provides a versatile and scalable approach for probing the RNA structurome. The experimental protocol requires minimal sample manipulation and can be directly applied to primary cells, and possibly animal tissues, enabling interrogation of endogenous RNAs in their native cellular context. Through integration with complementary techniques and future technical refinements, DMS-TRAM-seq has the potential to transform our understanding of how RNA structure shapes gene regulation. We anticipate that DMS-TRAM-seq will be instrumental in unraveling the structural regulatory code of the transcriptome, providing new opportunities to explore RNA-based mechanisms of gene regulation, disease pathology, and therapeutic intervention.

## Acknowledgements

We thank Silvi Rouskin, Christopher Burge, David Bartel, and members of the Jain lab for helpful discussions, and Andy Nutter-Upham and Scott McCallum for the development of the humanRNAmap web portal. This work was supported by grants from the NIH (R35GM151111), Chan Zuckerberg Initiative, and the David and Lucile Packard Foundation to A.J.

## Author Contributions

K.F. and A.J conceived this study, designed experiments, and interpreted results. K.F. performed all experiments and most data analysis. T.W. developed code for generating transcriptome-wide mutational profiles. A.C. developed the pipeline for secondary structure prediction based on the mutational profiles. K.F. and A.J. wrote the paper with contributions from all authors.

## EXPERIMENTAL MODEL DETAILS

### Cell culture

Human U-2OS cells were cultured using DMEM (Life Technologies, 11965126) supplemented with 10% (v/v) FBS (Gibco, A56707-01) and 1% (v/v) antibiotic (Gibco, 15240-062), incubated at 37°C and 5% (v/v) CO_2_.

## METHOD DETAILS

### Arsenite/DMS treatment and RNA isolation

Each sample of U-2OS cells was grown to ∼80% confluency in a single 10 cm plate. For cells treated with sodium arsenite (Sigma Aldrich, S7400-100G), sodium arsenite was added to the media at a final concentration of 0.5 mM for 30 minutes. During this time, cells continued to incubate at 37°C and 5% CO_2_. For cells treated with DMS, a separate 15 mL/sample solution of 2% DMS (Sigma Aldrich, D186309-5ML) in DMEM was created, using pre-warmed (37°C) DMEM that did not contain FBS or antibiotic. Vigorous shaking of the DMS in DMEM solution was required to ensure mixing of the DMS solution immediately before adding the solution to cells. The original growth media was removed from the cells, and the 15 mL 2% DMS solution was added to the plate for 3 minutes at room temperature.

The DMS solution was removed from the cells, and the cells were washed on-plate 3 times with 5 mL DPBS (Life Technologies, 14040133) each, as gently as possible to avoid cell detachment from the plate. 600 μL lysis buffer from the PureLink RNA Mini Kit (Life Technologies, 12183018A), containing 1% 2-mercaptoethanol (BME) (VWR, M131-250ML), was added directly to the plate, with the BME serving to quench any trace remaining DMS. As directed by the PureLink protocol, 600 μL 70% ethanol was added to each sample. Only at this point were samples removed from the safety hood. The rest of the PureLink RNA Mini Kit protocol was followed, and samples were eluted in 50 μL or 30 μL of nuclease-free water (Thermo Fisher Scientific, AM9932) for non-DMS and DMS-treated samples, respectively. Total RNA samples were run on a BioAnalyzer to ensure quality before ribodepletion.

### Ribodepletion

7 μL of total RNA was added to a PCR tube containing 1 μL of 10ug/μL custom ribodepletion oligos (Table S9) in 2 μL of 5x hybridization buffer (final concentrations: 100 mM NaCl and 100 mM Tris-HCl at pH 7.5 in nuclease-free water). Samples were transferred to a pre-heated thermocycler at 95°C for 2 minutes, then ramped to 45°C at -0.1°C/s. An RNase H solution consisting of 0.5 μL nuclease-free water, 1.5 μL Thermostable RNase H (VWR, 76081-726), and 3 μL 5x RNase H buffer (final concentrations: 10 mM MgCl2 and 33.3 mM NaCl) was then added directly to the hybridization mix at 45°C, and the samples were incubated at 45°C for 30 minutes. The RNA was then purified using 2.2x (33 μL) RNAClean XP beads (Beckman Coulter, A63987) per manufacturer’s protocol, and eluted in 16.5 μL nuclease-free water.

To digest the ribodepletion oligos and any genomic DNA in the sample, 1.5 μL Turbo DNase and 2 μL 10x Turbo DNase buffer were added to each sample, then incubated at 37°C for 30 minutes. Another 2.2x (44 μL) RNAClean XP cleanup was performed, and the ribodepleted RNA was eluted in 10 μL nuclease-free water.

### Fragmentation & Reverse Transcription

The fragmentation reagents and primers used for reverse transcription were from the IDT xGen Broad Range Library Prep Kit (IDT, 10009813), though using a custom protocol. 8 μL of ribodepleted RNA (recommended: >100 ng per sample) was added to a PCR tube containing 1 μL F1 reagent (random primers), 4 μL F3 reagent, and 2 μL F4 reagent from the IDT xGen Broad Range Library Prep Kit. Samples were placed in a pre-heated 95°C incubator for 0-2 minutes for fragmentation, depending on the RNA’s size distribution as observed via Bioanalyzer after ribodepletion, and then immediately placed on ice for primer annealing.

A reverse transcription mix, containing 2 μL TGIRT reverse transcriptase (InGex, TGIRT5x50), 1 μL RNase inhibitor (R1 from the xGen kit or RNaseOUT, Life Technologies 10777019), and 1 μL 0.1 M DTT was added directly to the fragmentation mixture, and the samples were incubated at room temperature for 30 minutes in order to enable binding of TGIRT to the RNA/primer duplex. After this incubation, 2 μL F2 reagent (dNTPs) were added to each reaction, and the samples were put in a preheated thermocycler with the following program: 20°C for 10 minutes, 42°C for 10 minutes, and 55°C for 1 hour. After this hour, 1 μL 4M NaOH was added to hydrolyze the RNA and to inactivate TGIRT. Then, the samples were heated to 95°C for 3 minutes before being cooled to a 4°C hold. Samples were then neutralized using 1 μL 4M HCl. 27 μL low EDTA TE buffer (from the xGen kit) was added to each sample, and then the cDNA was purified in a 1.0x (50 μL) Ampure XP bead cleanup and eluted in 10 μL low EDTA TE (from xGen Broad Range Library Prep Kit).

### Library prep & sequencing

After the cDNA is purified as described above, the standard protocol for the xGen Broad Range RNA Library Prep kit was followed. In the PCR step, unique dual indexing primers were used, and 10 PCR cycles were performed at thermocycler settings recommended by the xGen protocol. Two 0.85x Ampure XP bead cleanups were performed to thoroughly remove remaining primer. Samples were multiplexed and sequenced on Illumina NovaSeq 6000 using 150-base paired-end reads and 8 cycle reads at both indices, using the appropriate flow cell for an expected output of ∼150-200 million read pairs per sample.

## QUANTIFICATION AND STATISTICAL ANALYSIS

### Read processing and genome alignment

For quality and adapter trimming, *cutadapt* was used with the following options:

cutadapt -a AGATCGGAAGAGCACACGTCTGAACTCCAGTCA -A
AGATCGGAAGAGCGTCGTGTAGGGAAAGAGTGT -u 0 -U 10 -q 20 -j 12 -m 10

Next, the reads were deduplicated by sequence in order to remove PCR duplicates using *clumpify*, part of the BBMap package, with the following options: dedupe=t, optical=f, subs=1.

*STAR* was then used for alignment to the hg38 human reference genome at default settings, with the output option –outSAMtype BAM Unsorted.

*Samtools sort* was used to sort the bam files by coordinate, and all non-unique (unaligned and multimapping) reads were removed using *samtools view* with the -b, -F 260, and -q 20 options.

### Mutational profile calculation & filtering

In order to calculate the mismatch rates per reference base, *bcftools mpileup* was used with the default base alignment quality (BAQ) score setting, to ensure that mismatch rates are not affected by local misalignments, as well as these custom options:

- d 1000000 -q 20 -Q 20 -a format/ad,info/ad.

*Bcftools norm* and *bcftools query* were then used to manipulate the resulting variant call format (vcf) file, to generate a tab-separated table of base call counts at each reference position. This file was then further processed using a custom python script to calculate the overall coverage and mismatch rates per reference base (see process_pileup_strand_select.py).

These raw mutational profiles were filtered by coverage, where only reference bases with at least 100 aligned reads per base were retained for each sample. Then, *bedtools intersect* was used to only retain bases meeting this threshold across all DMS and control samples. For each DMS-treated sample, the mismatch rates for the corresponding non-DMS-treated replicate were subtracted in order to correct for SNPs and any endogenous modifications that may be read out as mismatches without DMS treatment. The resulting “normalized” mutational profile was capped with a lower value of 0, to account for any negative values from the normalization. This coverage-filtered and control-corrected dataset was the final mutational profile used in all further described analyses, and is available on the Zenodo repository.

### Annotation-based analyses

This analysis is designed to use an input bed annotation file to define the regions of interest. For the coordinates defined by each row in the bed file, *bedtools intersect* and *bedtools groupby* were used to collect and group the relevant mismatch rates for the region, resulting in a list of mismatch rates connected to each bed file row. This data was then processed by a custom python script, process_replicate_signal.py, which calculated several metrics of interest, including the average mismatch rate, Gini index, and Pearson’s R. In experiments with multiple conditions, comparative statistics were also performed on each of those metrics, yielding fold-change and p-values.

### Window-based analyses

For each canonically-defined transcript, windows of a set size (i.e. a defined number of A/C datapoints passing all coverage filters, not actual sequence length on the transcript) were tiled across the genome. For each window, a number of statistics were calculated. To identify structured regions within one condition, the Gini index was calculated for each replicate, as well as the average Pearson’s correlation coefficient in pairwise replicate comparisons.

To identify local changes in structure, additional metrics were calculated, including the quantity δ, defined as:

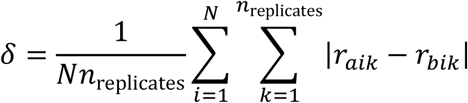

Where *i* is an index of A/C nucleotides within the window, N is number of A/C nucleotides within the window, k is an index for the number of replicates, r_a_ is the mismatch rate in condition a, and r_b_ is the mismatch rate in condition b.

To calculate the significance of structural change within these windows, we calculated a p-value comparing two sets of Pearson’s R values. The first set of R values, R_within_, was calculated by comparing the mismatch rates between replicates of the same condition, in order to understand the general signal reliability of the region in question. The second set of R values, R_between_, was calculated by comparing the mismatch rates between samples in the two different conditions. R_within_ and R_between_ are then compared using an independent t-test. Should the R values be comparable, resulting in a non-significant p-value, any difference in signal resulting from a difference in condition is indistinguishable from a difference in signal resulting from random replicate-to-replicate variability. Significant p-values, however, indicate a distinguishable difference between replicate-to-replicate variability and changes in signal resulting from the applied perturbation. P-values were corrected using the Benjamini-Hochberg method.

### Structure prediction & scoring

A region of interest for secondary structure prediction is defined by inputting the chromosome, coordinates, and strandedness of the gene into our custom structure prediction pipeline. Based on these values, the relevant DMS-TRAM-seq signal, as well as the relevant reference sequence, is pulled from the full dataset. The mismatch rate values are winsorized with only a top 5% limit (0%, 95% limits) to remove outliers, and then the values are scaled to a maximum of 1. These scaled values are used as constraints in the RNAstructure Fold --SHAPE option. The resulting structure output was input into VARNA for final image generation and overlay of the DMS-TRAM-seq signal colormap.

To analyze the accuracy of a given structure prediction (or the accuracy of DMS-TRAM-seq signal relative to a known control structure, as in Figure 2), we then calculated ROC curves utilizing sklearn.metrics.roc_curve and sklearn.metrics.roc_auc_score in python. “Positive” values are bases predicted to be unpaired; therefore, “true positives” are bases with high DMS-TRAM-seq signal that are also predicted to be unpaired, and “false positives” are bases with high DMS-TRAM-seq signal but are predicted to be paired.

### eCLIP & RBP analyses

20 A/C datapoint windows were calculated throughout the transcriptome, with several metrics of interest calculated for each window, as described above. In addition, the locations of each window were compared to a compilation of known CLIP binding sites for over 150 RBPs, where labels for the respective RBP binding sites were applied to the windows using the BEDOPS bedmap tool.

For each RBP, the total number of windows containing known binding sites (“target windows”) was determined, as well as the number of target windows significantly changing in accessibility, determined as discussed above (p_adj_ ≤ 0.05). In addition, a null model was made by scrambling the locations of all RBP binding sites. This randomization only occurred within windows known to have any annotated CLIP sites; windows with no known CLIP peaks were not included in this randomization. For this null model, the number of significantly-changing “target” windows was calculated. For each RBP, the ratio of the number of significantly-changing target windows in the data versus the null model yielded the “fold change” value. A positive fold change corresponds to a situation where the actual data contains more significantly-changing windows than the null model, and vice versa.

Next, for each RBP, a p-value is calculated using the hypergeometric test, with the following value inputs (as defined in the hypergeom.pmf function in scipy.stats, in python): k = significantly changing RBP target windows; M = total number of windows in entire dataset; n = the number of significantly changing windows in the entire dataset; N = the number of total RBP target windows. These statistics were calculated for all RBPs, and to select the best hits for display in Figure 5D, those with a log_2_FC of greater than 1.1 (or less than -1.1) and with p-values less than 0.001 were selected.

## ADDITIONAL RESOURCES

Online database for transcriptome-wide RNA secondary structure prediction based on DMS-TRAM-seq data: humanRNAmap.wi.mit.edu

## RESOURCE AVAILABILITY

### Lead contact

Requests for further information and resources should be directed to and will be fulfilled by the lead contact, Ankur Jain.

### Materials availability

This study did not generate new unique reagents.

### Data and code availability

Raw sequencing (FASTQ) data have been deposited at Sequence Read Archive at BioProject: PRJNA1246337.

Mutational profiling data and all original code have been deposited at Zenodo and are available at DOI 10.5281/zenodo.

