## Supplemental Information for "Transcriptome-wide RNA accessibility mapping reveals structured RNA elements and pervasive conformational rearrangements under stress"

Figure S1

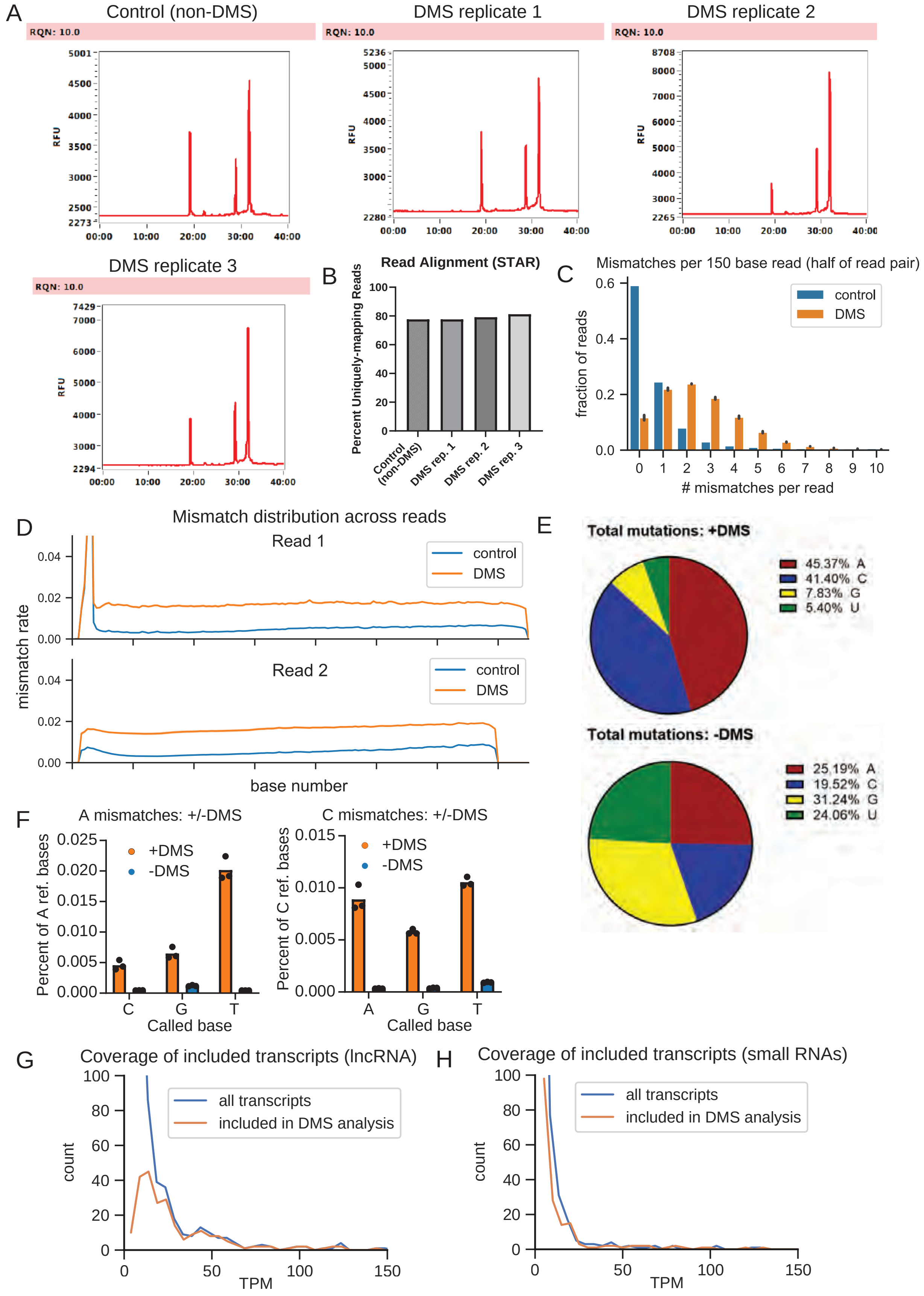
A
Control (non-DMS)
DMS replicate 1
DMS replicate 2
DMS replicate 3
RQN: 10.0
RFU
B
Read Alignment (STAR)
Percent Uniquely-mapping Reads
Control (non-DMS)
DMS rep. 1
DMS rep. 2
DMS rep. 3
C
Mismatches per 150 base read (half of read pair)
fraction of reads
control
DMS
### mismatches per read
D
Mismatch distribution across reads
Read 1
Read 2
mismatch rate
base number
E
Total mutations: +DMS
45.37% A
41.40% C
7.83% G
5.40% U
Total mutations: -DMS
25.19% A
19.52% C
31.24% G
24.06% U
F
A mismatches: +/-DMS
C mismatches: +/-DMS
+DMS
-DMS
Percent of A ref. bases
Percent of C ref. bases
Called base
G
Coverage of included transcripts (lncRNA)
H
Coverage of included transcripts (small RNAs)
all transcripts
included in DMS analysis
count
TPM

**Figure S1**. DMS-TRAM-seq library prep and genomic alignment. Related to Figures 1 and 2. (A) Bioanalyzer traces of total RNA from a non-DMS sample and three DMS-treated replicates, with RNA quality numbers (RQNs) labeled in the red boxes. (B) Percent of reads uniquely aligning to the genome, aligned with the STAR read aligner using default parameters (see Methods), for a non-DMS control sample and three DMS-treated replicates as in (A). (C) Mismatch frequency across each 150-base read end (half of a total read pair) in both a non-DMS control sample and averaged DMS-treated replicates (n = 3). (D) Distribution of mismatches across the two reads, for both a non-DMS control sample and the average of three DMS-treated replicates. Read 2 is 10 nt shorter than read 1 due to trimming of the length of the random primers. The peak at the beginning of read 1 is due to a small poly-G tail added during the annealing step of the library prep, and is consistent across all DMS and non-DMS samples. (E) Mutation frequency by reference base. For negatively-stranded genes, the reference bases were corrected to reflect the transcript's sequence. (F) Mutation rates for all possible called bases at reference A(left) and C (right) positions. (G) As in Figure 2A (main text), the distribution of TPM values for all annotated lncRNAs (blue) compared to lncRNAs meeting inclusion thresholds for further DMS-TRAM-seq analysis and structure prediction (orange). (H) Same as (G), but for small noncoding RNAs (snoRNAs, scaRNAs, snRNAs, scRNAs, and miRNAs).

**Table S1**: Transcript types included in DMS-TRAM-seq analysis, as annotated by UCSC genome browser. Related to Figure 2. See Table S2 for full characterization of each transcript.

| Gene type | Count |
|---|---|
| protein-coding | 8373 |
| lncRNA | 265 |
| pseudogenes | 164 |
| snoRNA | 100 |
| miRNA | 41 |
| misc. RNA (noncoding) | 31 |
| mt tRNA | 17 |
| snRNA | 12 |
| other (TEC, IG V) | 9 |
| scaRNA | 7 |
| ribozyme | 2 |
| scRNA | 1 |

Figure S2

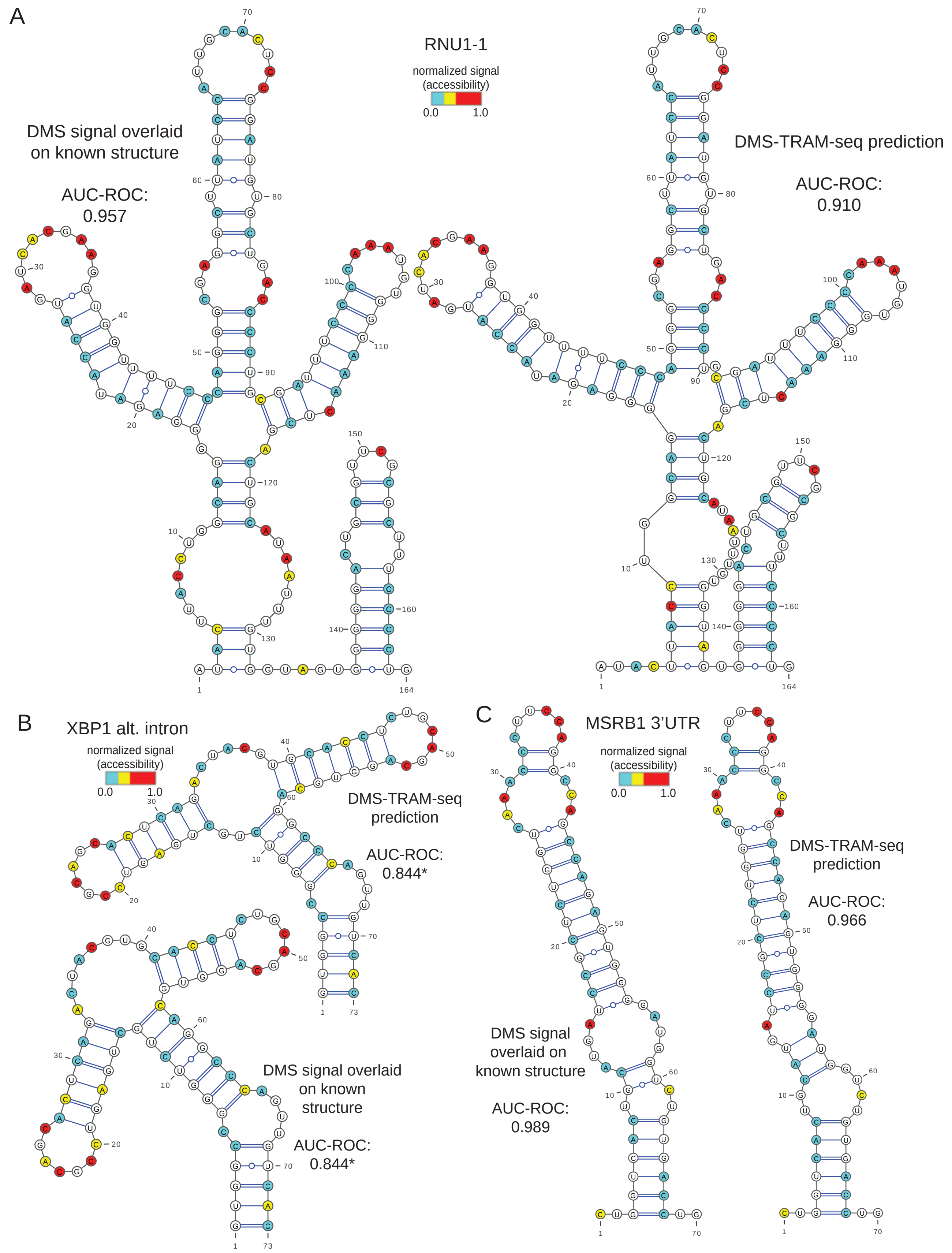
A
RNU1-1
normalized signal
(accessibility)
0.0
1.0
DMS signal overlaid
on known structure
AUC-ROC:
0.957
DMS-TRAM-seq prediction
AUC-ROC:
0.910
B
XBP1 alt. intron
normalized signal
(accessibility)
0.0
1.0
DMS-TRAM-seq
prediction
AUC-ROC:
0.844*
DMS signal overlaid
on known
structure
AUC-ROC:
0.844*
C
MSRB1 3'UTR
normalized signal
(accessibility)
0.0
1.0
DMS-TRAM-seq
prediction
AUC-ROC:
0.966
DMS signal
overlaid on
known structure
AUC-ROC:
0.989

**Figure S2**. Control structures for DMS-TRAM-seq benchmarking. Related to Figure 2. (A) Left, DMS-TRAM-seq signal overlaid as a colormap (blue = low, yellow = moderate, red = high accessibility) on the published structure of *RNU1-1* (Tomezsko et al., 2020)[37]. Right, structure prediction of *RNU1-1* constrained by DMS-TRAM-seq signal, with an identical colormap. AUC-ROC values quantify agreement between DMS-TRAM-seq signal and each structure. (B) Same as (A), but for a known structure in the *XBP1* alternate intron (Rouskin et al., 2014)[44]. (C) Same as (A), but for a known structure in the 3'UTR of *MSRB1* (Rouskin et al., 2014)[44].

Figure S3

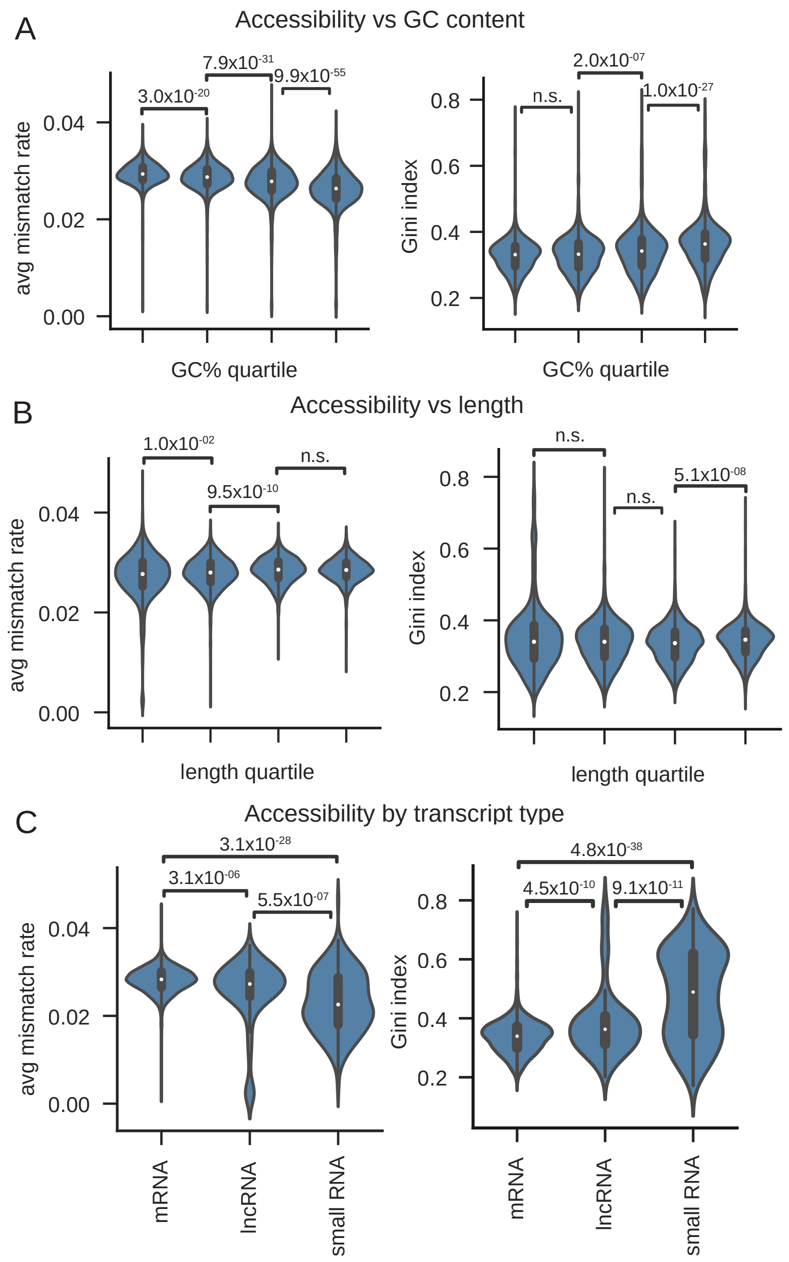
A
Accessibility vs GC content
3.0x10-20
7.9x10-31
9.9x10-55
avg mismatch rate
0.04
0.02
0.00
GC% quartile
n.s.
2.0x10-07
1.0x10-27
Gini index
0.8
0.6
0.4
0.2
GC% quartile
B
Accessibility vs length
1.0x10-02
9.5x10-10
n.s.
avg mismatch rate
0.04
0.02
0.00
length quartile
n.s.
n.s.
5.1x10-08
Gini index
0.8
0.6
0.4
0.2
length quartile
C
Accessibility by transcript type
3.1x10-28
3.1x10-06
5.5x10-07
avg mismatch rate
0.04
0.02
0.00
mRNA
lncRNA
small RNA
4.8x10-38
4.5x10-10
9.1x10-11
Gini index
0.8
0.6
0.4
0.2
mRNA
lncRNA
small RNA

**Figure S3**. Global trends in RNA accessibility. Related to Figure 3. (A) Distributions of per-transcript average mismatch rates (left) and Gini indices (right), stratified by GC-content quartile (low to high). (B) Same as (A), stratified by transcript length quartile (low to high). (C) Same as (A), but transcripts are grouped by type. Small RNAs include snoRNAs, scaRNAs, snRNAs, scRNAs, and miRNAs.

**Table S3**: Transcript types within highly-structured transcripts (R ≥ 0.6, Gini ≥ 0.4). Related to Figure 3.

| **Gene type** | **Count** |
|---|---|
| protein-coding | 163 |
| snoRNA | 42 |
| lncRNA | 19 |
| snRNA | 11 |
| misc. RNA (noncoding) | 7 |
| pseudogenes | 6 |
| scaRNA | 5 |
| miRNA | 3 |
| ribozyme | 2 |
| scRNA | 1 |

Figure S4

A

*S. cerevisiae* RNase MRP RNA
cryo-EM structure
Perederina et al., Nature Communications 2020

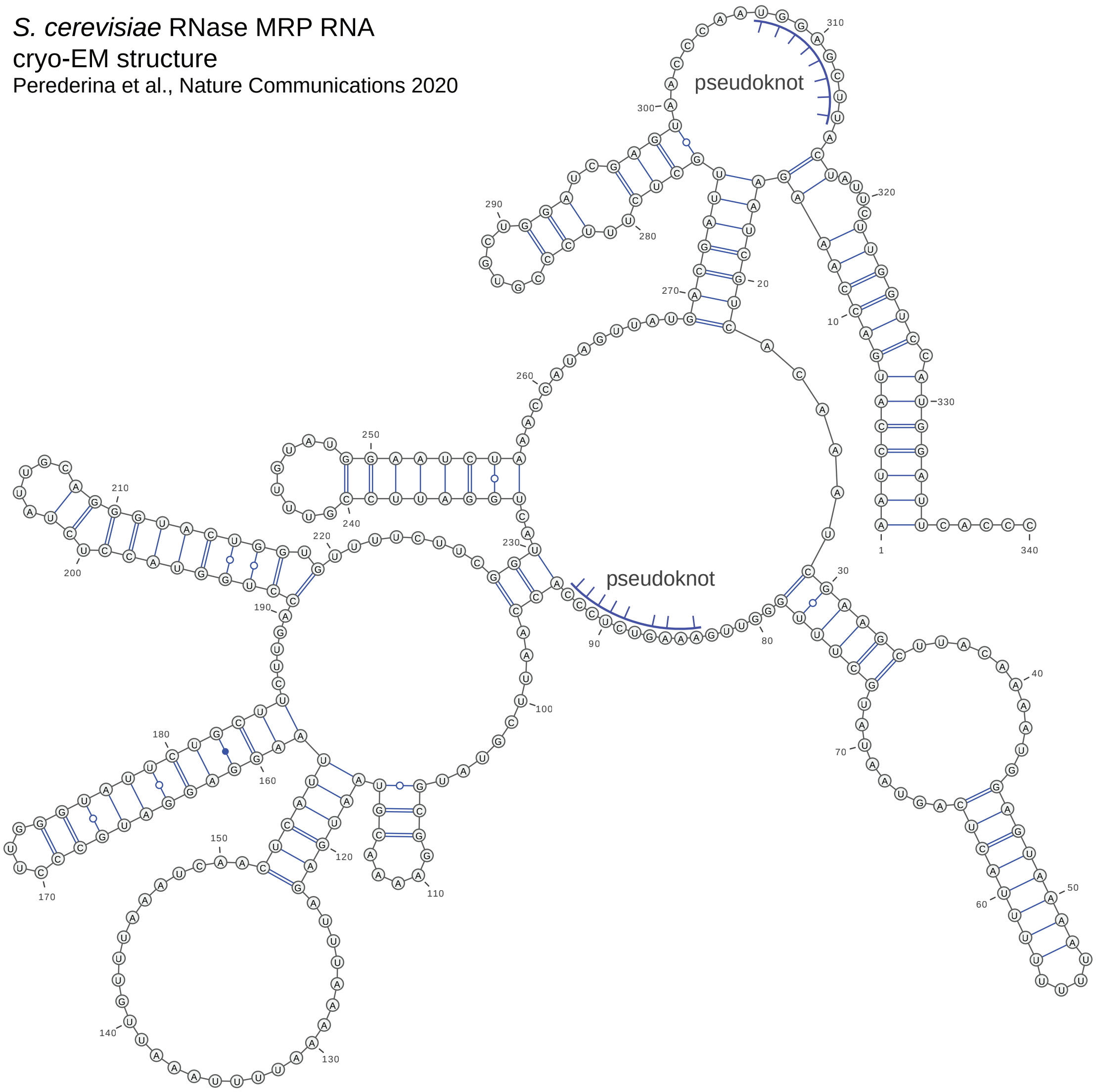


B

R-scape/R2R output: RMRP eukaryotic alignment

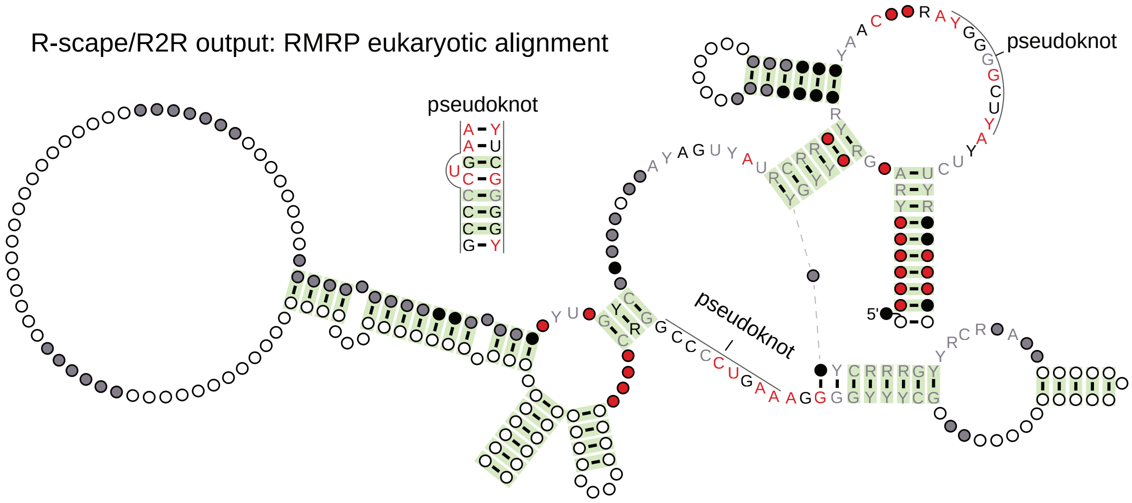

**Figure S4**. Yeast *RMRP* structure & covariation analysis. Related to Figure 3. (A) Secondary structure of *S. cerevisiae RMRP* (adapted from Perederina et al., 2020)[51]. (B) Evolutionary covariation analysis of 933 eukaryotic homologs (R-scape; Rivas et al., 2017)[57]. Significantly covarying base pairs ($E \leq 0.05$) highlighted in green (see Methods).

Figure S5

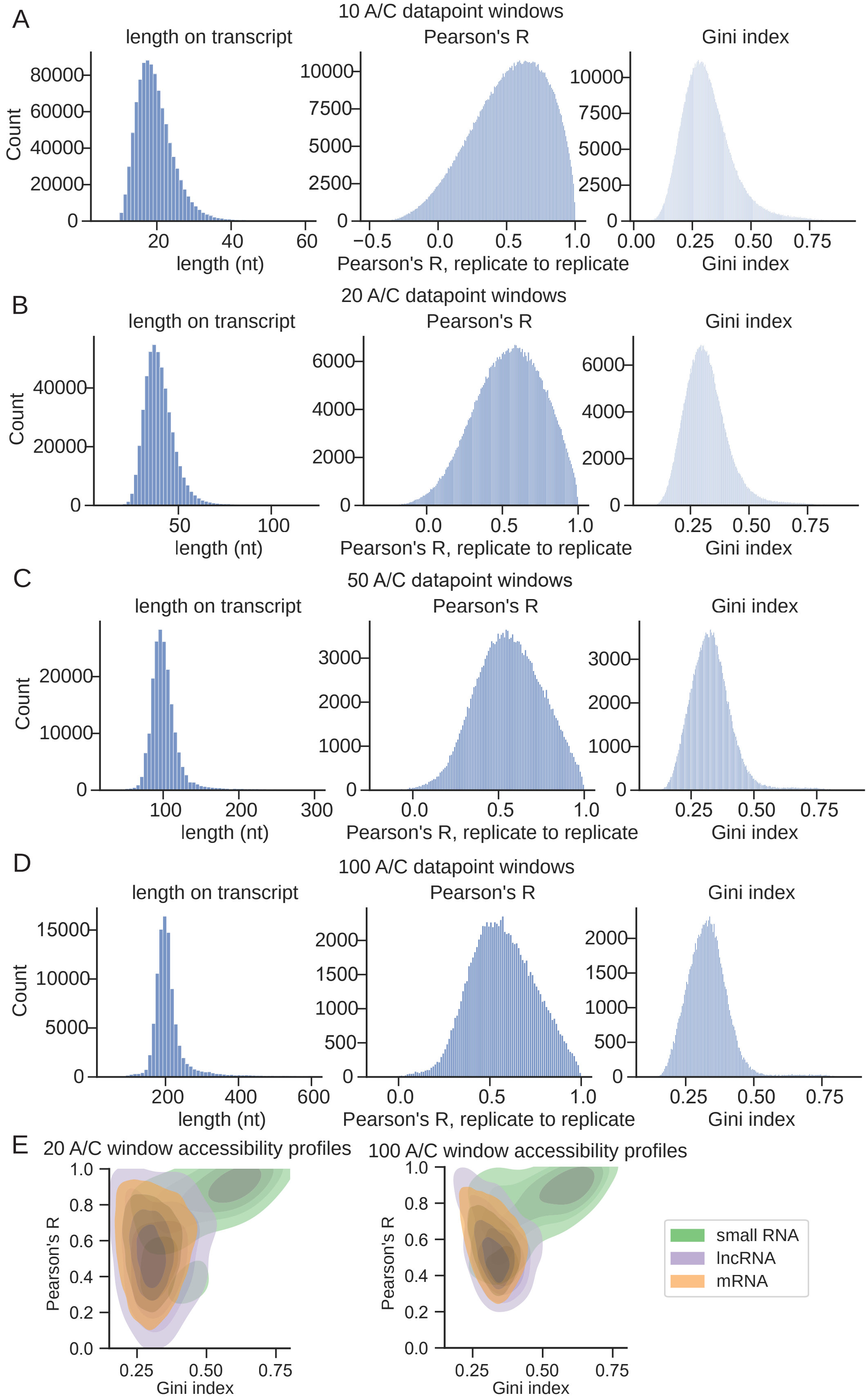
A
10 A/C datapoint windows
length on transcript
Pearson's R
Gini index
Count
length (nt)
Pearson's R, replicate to replicate
Gini index
B
20 A/C datapoint windows
length on transcript
Pearson's R
Gini index
Count
length (nt)
Pearson's R, replicate to replicate
Gini index
C
50 A/C datapoint windows
length on transcript
Pearson's R
Gini index
Count
length (nt)
Pearson's R, replicate to replicate
Gini index
D
100 A/C datapoint windows
length on transcript
Pearson's R
Gini index
Count
length (nt)
Pearson's R, replicate to replicate
Gini index
E
20 A/C window accessibility profiles
100 A/C window accessibility profiles
Pearson's R
Gini index
small RNA
lncRNA
mRNA

**Figure S5**. Characterization of various window sizes. Related to Figure 3. (A) Distributions of several metrics for 10 A/C datapoint windows: sequence lengths (left), inter-replicate Pearson's correlation coefficients (middle), and Gini indices (right). (B) Same as (A), but for 20 A/C datapoint windows, as used in Figure 6 for RBP analysis. (C) Same as (A), but for 50 A/C datapoint windows, as used for the detection of structured elements (Figure 3) and structural changes (Figure 5). (D) Same as (A), but for 100 A/C datapoint windows. (E) Joint distribution of Gini index versus Pearson's R for 20 (left) and 100 (right) A/C datapoint windows.

Figure S6

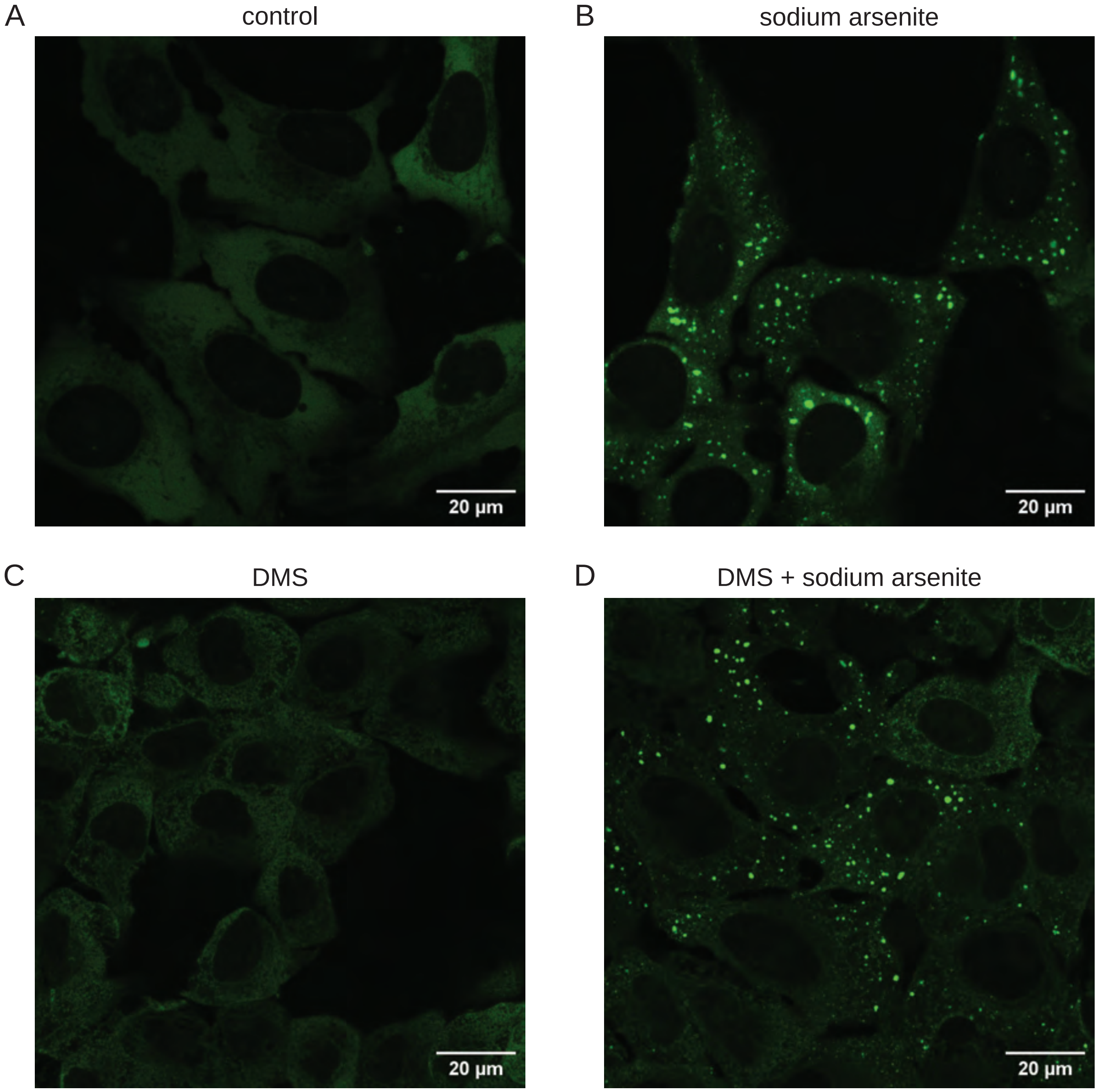
A
control
20 µm
B
sodium arsenite
20 µm
C
DMS
20 µm
D
DMS + sodium arsenite
20 µm

**Figure S6**. Stress granule formation in arsenite- and DMS-treated cells expressing G3BP1-GFP. Related to Figures 4-6. Cells were fixed immediately after respective treatments with 2% formaldehyde. (A) G3BP1-GFP signal in cells without stress applied. (B) G3BP1-GFP signal in cells treated with 0.5 mM sodium arsenite for 30 minutes. (C) G3BP1-GFP signal in cells treated with 2% DMS for 3 minutes. (D) G3BP1-GFP signal in cells treated with 0.5 mM sodium arsenite for 30 minutes, followed by 2% DMS for 3 minutes.

Figure S7

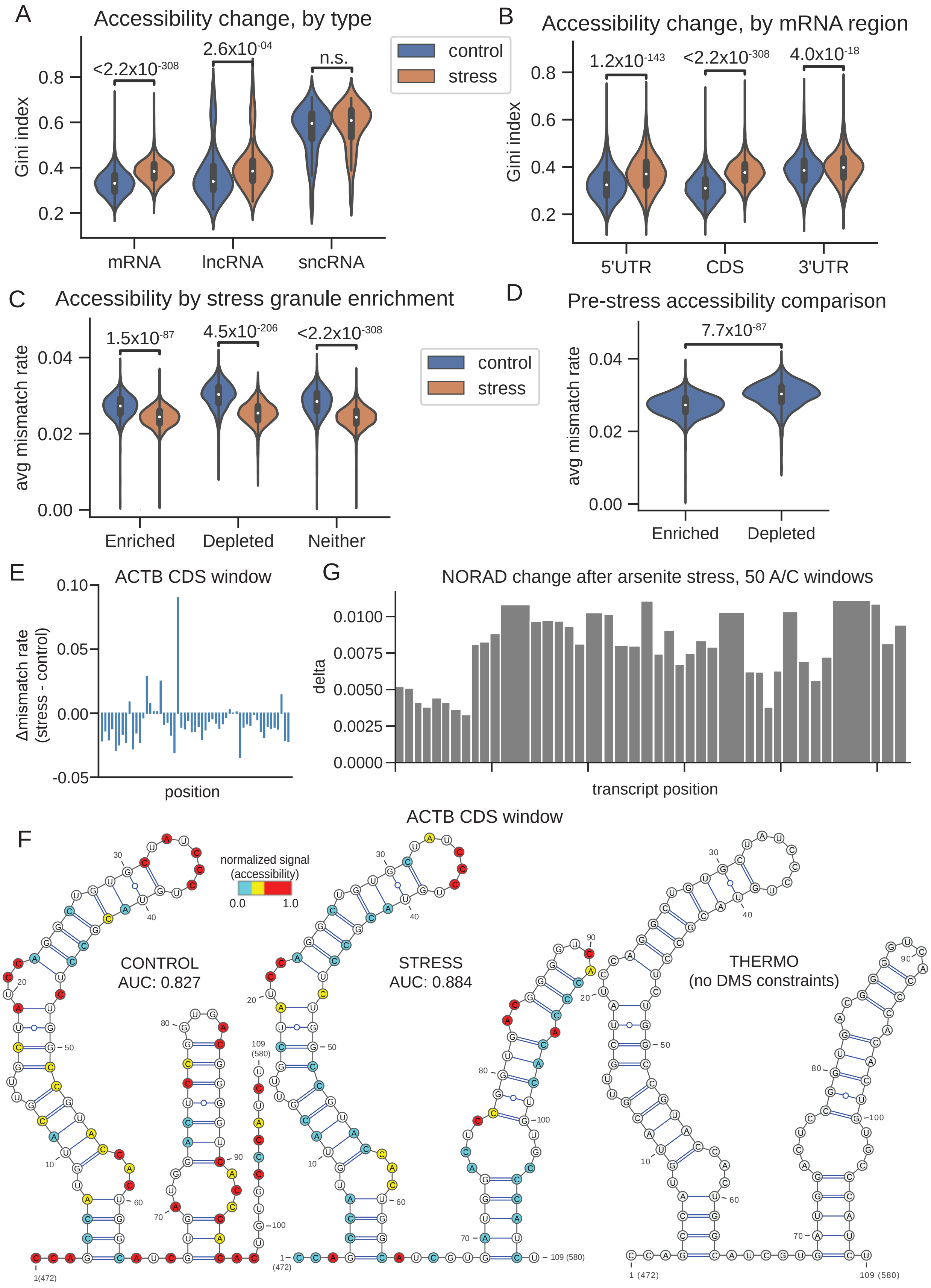

A
Accessibility change, by type
Gini index
<2.2x10-308
2.6x10-04
n.s.
mRNA
lncRNA
sncRNA
control
stress
B
Accessibility change, by mRNA region
Gini index
1.2x10-143
<2.2x10-308
4.0x10-18
5'UTR
CDS
3'UTR
C
Accessibility by stress granule enrichment
avg mismatch rate
1.5x10-87
4.5x10-206
<2.2x10-308
Enriched
Depleted
Neither
control
stress
D
Pre-stress accessibility comparison
avg mismatch rate
7.7x10-87
Enriched
Depleted
E
ACTB CDS window
Δmismatch rate (stress - control)
position
G
NORAD change after arsenite stress, 50 A/C windows
delta
transcript position
ACTB CDS window
F
normalized signal (accessibility)
CONTROL
AUC: 0.827
STRESS
AUC: 0.884
THERMO
(no DMS constraints)

**Figure S7**. DMS-TRAM-seq signal changes after arsenite stress. Related to Figures 4 and 5. (A) Gini index distributions across transcripts, grouped by transcript type, comparing untreated (blue) and arsenite-treated (orange) conditions. (B) Same as (A), but grouped by mRNA subregion. (C) Average mismatch rate distribution for transcripts found to be enriched in or depleted from stress granules, as reported in Khong et al., 2017[72]. (D) Average mismatch rate distribution comparing the accessibilities of stress-granule-enriched and stress-granule-depleted transcripts in untreated cells. (E) For a 50 A/C datapoint window within the CDS of ACTB, the changes in mismatch rate (stress – control) per base. This region spans positions 472 to 580 on the canonical transcript. (F) For the same window as in (E), the DMS-TRAM-seq constrained structure prediction in the control condition (left), stress condition (center), and the prediction based on sequence alone, without any constraints applied (right). (G) For all 50 A/C datapoint windows in NORAD, the $\delta$ value (average bidirectional change per base after arsenite stress) is given. Windows in the 5' region of the transcript are observed to change less than those in the rest of the transcript, as observed in prior publications (Farberov et al., 2024)[42]. P-values calculated via two-sided Mann-Whitney U test.

Figure S8

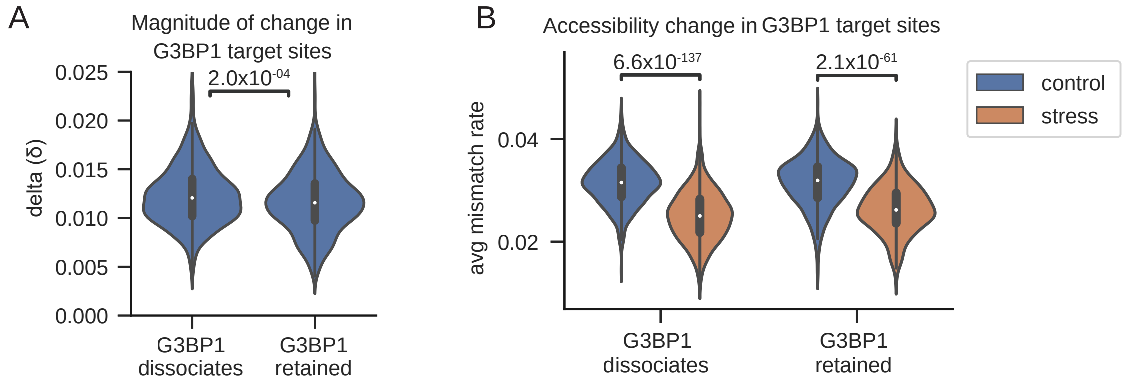
A
Magnitude of change in
G3BP1 target sites
2.0x10-04
delta (δ)
0.025
0.020
0.015
0.010
0.005
0.000
G3BP1
dissociates
G3BP1
retained
B
Accessibility change in G3BP1 target sites
6.6x10-137
2.1x10-61
avg mismatch rate
0.04
0.02
G3BP1
dissociates
G3BP1
retained
control
stress

**Figure S8**. RBP-dependent RNA structural changes during arsenite stress revealed by DMS-TRAM-seq. Related to Figure 6. (A) Distribution of δ values (average bidirectional change per base after arsenite stress) for all 20 A/C datapoint windows where G3BP1 is known to dissociate or be retained throughout the arsenite stress response (Xiao et al., 2024)[77]. (B) Distributions of average mismatch rates across G3BP1-dissociating versus G3BP1-retained sites in control (blue) and stress (orange) conditions.

**SUPPLEMENTAL INFORMATION INDEX**

Table S2. Full transcript-level results and characterization. Related to Figure 2 and 3A-B. Table S4. Full dataset of 50-datapoint windows, related to Figure 3.

Table S5. Table with full metrics for all 721 highly-structured elements, as shown in Figure 3.

Table S6. All 50-datapoint windows from the arsenite vs control comparative analysis, as shown in Figure 5.

Table S7. All 20-nucleotide windows from the arsenite vs control comparative analysis, including cross-referenced eCLIP peaks, related to Figure 6.

Table S8. Full RBP-level analysis results, related to Figure 6.

Table S9. Sequences of ribodepletion oligonucleotides, as described in Methods.
